# Lung-Selective Immune Reprogramming via *In Situ* Red Blood Cell Hitchhiking Nanoparticles

**DOI:** 10.64898/2026.02.27.708550

**Authors:** Edidiong M. Udofa, Endong Zhang, Mohammad Moein Naderi, Shan He, Hirushi Gunasekara, Bailin Feng, Chih-Jia Chao, Luyu Zhang, Jeonghun Lee, Petra Čechová, Markéta Paloncýová, Dejan S. Nikolic, Margarita Herrera-Alonso, Ying Liu, Ying S. Hu, Zhangli Peng, Michal Otyepka, Zongmin Zhao

**Affiliations:** Department of Pharmaceutical Sciences, University of Illinois Chicago, Chicago, Illinois, USA; Department of Biomedical Engineering, University of Illinois Chicago, Chicago, Illinois, USA; Department of Chemistry, University of Illinois Chicago, Chicago, Illinois, USA; Department of Chemical Engineering, University of Illinois Chicago, Chicago, Illinois, USA; School of Materials Science and Engineering, Colorado State University, Fort Collins, Colorado, USA; Regional Center of Advanced Technologies and Materials, The Czech Advanced Technology and Research Institute (CATRIN), Palacký University Olomouc, Czech Republic; UICentre for Drug Discovery, University of Illinois Chicago, Chicago, Illinois, USA; School of Biomedical and Chemical Engineering, Colorado State University, Fort Collins, Colorado, USA; University of Illinois Cancer Center, Chicago, Illinois, USA; IT4Innovations, VŠB – Technical University of Ostrava, Ostrava-Poruba, Czech Republic

## Abstract

Premature clearance and limited organ targeting remain major barriers for nanoparticle (NP) drug delivery. Hitchhiking NPs on red blood cells (RBCs) can enhance circulation and organ-selective accumulation, but most approaches require *ex vivo* RBC extraction and reinfusion, limiting clinical translation. Here, we report an *in situ* RBC-hitchhiking strategy, named i-Bind, which employs polyphenol surface functionalization to enable spontaneous NP attachment to RBCs directly in the bloodstream. Driven by strong interactions of phenolic motifs with RBC membranes, i-Bind NPs exhibited markedly enhanced and more stable hitchhiking onto RBCs under flowing whole blood conditions. In both healthy and diseased mice, i-Bind NPs preferentially target the lungs, resulting in an over 20-fold increase in lung-to-liver deposition ratio compared to unmodified NPs. Additionally, i-Bind NPs show preferential targeting to distinct lung immune cell subsets in a pathology-dependent manner, including cDC2s in healthy lungs, neutrophils in acute lung injury, and cDC1s in lung metastases. In a melanoma lung metastasis model, delivery of the STING agonist diABZI via i-Bind NPs significantly inhibited lung metastasis progression by reprogramming the lung immune microenvironment. Collectively, i-Bind provides a simple and versatile platform for organ-targeted drug delivery and immune reprogramming.

**Teaser:** Surface functionalization of nanoparticles enables *in situ* red blood cell hitchhiking, unlocking new paths for organ-selective immune reprogramming

## Introduction

Nanoparticles (NPs) have been extensively investigated over the past few decades to improve drug pharmacological profiles, enable controlled release, prolong circulation, and facilitate targeted delivery(*1–3*). Despite encouraging clinical advances, conventional NPs continue to face long-standing challenges that limit their broader applications. Particularly, rapid clearance by the mononuclear phagocyte system (MPS), nonspecific accumulation in the liver and spleen, and limited deposition in target tissues often result in suboptimal therapeutic efficacy and increased systemic toxicity(*1, 3, 4*). These limitations underscore the need for new strategies that extend beyond optimizing synthetic carriers and instead exploit biological systems that have naturally evolved efficient trafficking and targeting mechanisms. Cellular hitchhiking, engineering living cells to enhance NP delivery, has emerged as a particularly promising approach in this context(*5–11*). By attaching NPs to circulating cells, NPs can leverage the natural distribution and homing patterns of living cells to achieve site-specific drug delivery(*12–14*). This strategy integrates the strengths of synthetic materials with biological carriers to improve therapeutic efficacy and minimize off-target toxicity. A particular notable example is red blood cell (RBC) hitchhiking, which leverages the unique biophysical properties of RBCs to deliver surface-bound NPs to selected organs such as the lungs(*12, 15, 16*). As RBCs continuously transit through microvasculature, their passage through narrow capillaries creates a unique biophysical opportunity: NPs adhered to their surface can be mechanically dislodged and deposited into the lungs(*12, 17, 18*). Using this strategy, up to 40% of intravenously administered NPs can accumulate in the lungs, compared to less than 5% with free NPs(*17*). RBC hitchhiking has demonstrated potential for improving the delivery of chemotherapeutics(*19*), immunotherapy(*16*), anti-inflammatory agents(*20*), antiviral agents(*21*), and gene therapies(*22, 23*) for treating pulmonary diseases.

However, despite these encouraging advances, the clinical translation of early RBC hitchhiking methods has been limited by their reliance on *ex vivo* cell modification(*7*). In these first-generation methods, RBCs are extracted, engineered with NPs, and reinfused. While effective as a proof-of-concept, this process is largely restricted by the availability and quality of RBC sources and is difficult to standardize. Additionally, *ex vivo* manipulation may compromise RBC membrane integrity, reducing cell deformability and lifespan, which are critical for maintaining effective circulation(*7, 24*). Moreover, *ex vivo* manipulations may elevate risks of immunogenicity, as modified RBCs may be recognized as foreign by the immune system, leading to rapid clearance and potentially triggering harmful immune responses(*7, 24*). These drawbacks underscore the need for next-generation strategies that enable efficient and stable NP-RBC interactions directly *in situ*, thereby avoiding the pitfalls of *ex vivo* manipulation and improving clinical feasibility(*18, 25*).

Here, we report an *in situ* RBC-hitchhiking NP platform, referred to as i-Bind NPs, which employs polyphenol surface functionalization to enhance spontaneous NP attachment to RBCs *in situ* directly in the bloodstream (**Fig. 1a**). i-Bind is based on coating polymeric NPs with tannic acid (TA), which forms tight and stable non-covalent interactions with diverse molecular partners on RBC membranes through hydrogen bonding, electrostatic interactions, hydrophobic interactions, and π–π stacking(*26–28*). Polyphenols have gained significant attention in biomaterial design because of their unique chemical structures and multifunctional properties(*27–30*). TA, in particular, possesses abundant phenolic groups that can engage in hydrogen bonding, electrostatic and hydrophobic interactions, and metal coordination(*31*). These versatile binding modes allow polyphenols to form strong, multivalent interactions with biomolecules, and they have been widely applied in the design of coatings, adhesives, and drug delivery systems(*29, 32*). By harnessing these properties of TA coating, i-Bind NPs provide a simple yet powerful strategy to engineer NP surfaces for efficient interactions with cellular membranes. Compared with other RBC-targeting strategies, such as antibody- or peptide-decorated NPs(*33, 34*), i-Bind leverages tannic acid-mediated RBC adhesion to achieve broad, reproducible, and disease-adaptive cargo delivery, while avoiding the complexity, variability, and immunogenic risks inherent to other targeted approaches. We show that the i-Bind NPs achieved efficient *in situ* hitchhiking onto RBCs under physiological flow conditions. Following intravenous administration, i-Bind NPs demonstrated preferential accumulation in the lungs, and significantly reduced accumulation to the liver, resulting in a >20-fold increase in lung-to-liver deposition ratio compared to unmodified polymeric NPs. Additionally, beyond enhanced lung targeting, i-Bind NPs also demonstrated pathology-dependent immune cell accumulation, enabling distinct cell-level biodistribution across different disease states. Leveraging this enhanced pulmonary delivery, we demonstrated that delivery of the STING agonist dimeric amidobenzimidazole (diABZI) using i-Bind NPs significantly inhibited metastatic progression in a B16F10 melanoma lung metastasis model, consistent with modulation of the local immune microenvironment, enhancing cDC1 recruitment and activation, and increasing CD8⁺ T cell infiltration. Collectively, by enabling efficient *in situ* RBC hitchhiking through polyphenol surface functionalization, i-Bind potentially overcomes the translational barriers of *ex vivo* RBC modification. Our findings suggest that i-Bind can serve as a simple and versatile platform for organ- and immune cell-preferential drug delivery.

**Fig. 1.**
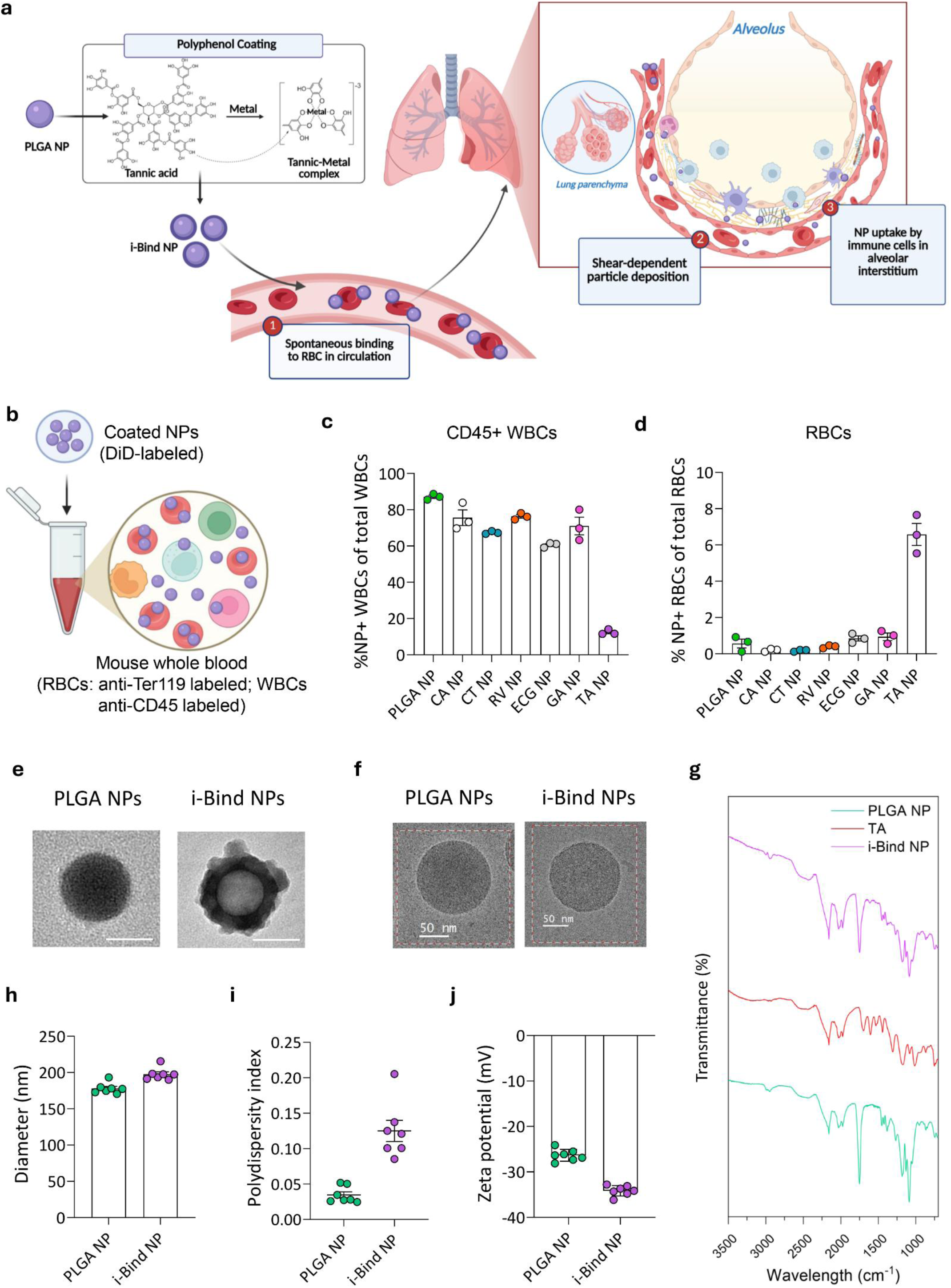
Design and characterization of i-Bind NPs for *in situ* RBC hitchhiking. **a,** Schematic depicting the design of i-Bind NPs for *in situ* RBC binding and lung delivery. Created in BioRender. https://BioRender.com/rwpnw3u. **b,** Schematic showing the experimental setup to screen polyphenol-coated NPs for RBC binding in whole blood. Created in BioRender. Zhao, Z. (2025) https://BioRender.com/76otyy0. **c-d,** Binding of polyphenol-coated NPs to CD45^+^ WBCs **(c)** and RBCs **(d)** in whole blood. Percentage of NP-positive cells of total respective cells was shown. CA: caffeic acid; CT: catechin; RV: resveratrol; ECG: epigallocatechin gallate; GA: gallic acid; TA: tannic acid. **e-f,** Representative TEM images **(e)** and cryo-TEM images **(f)** of PLGA NPs and TA-coated (i-Bind) NPs. Scale bar: in (e), 100 nm; in (f), 50 nm. **g,** FTIR spectrum for i-Bind NPs showing TA coating onto PLGA NPs. **h-j,** Physicochemical properties of NPs, including hydrodynamic size **(h)**, polydispersity index **(i)**, and surface charge **(j)** of PLGA NPs and i-Bind NPs. Data in **(c, d, h-j)** are presented as mean ± SEM.

## Results

### Screening and characterization of i-Bind NPs for optimal RBC binding

We synthesized i-Bind NPs by coating the surface of poly(lactic-co-glycolic acid) (PLGA) NPs with a metal-phenolic network composed of phenolic compounds and metal ions such as iron (Fe^3+^). Fe^3+^ serves as a crosslinker and forms coordination complexes with multiple phenol molecules, resulting in the formation of a metal-phenolic network on the surface of PLGA NPs(*28, 35*). We prepared a set of NPs coated with different phenolic materials (**Table S1**) and screened them for optimal binding to RBCs in mouse whole blood. Each coated NP formulation was incubated with mouse whole blood, and the binding of NPs to RBCs and CD45^+^ white blood cells (WBCs) was quantified (**Fig. 1b**). All the tested phenol-coated NPs showed significantly reduced binding to CD45^+^ WBCs compared to uncoated NPs, with tannic acid (TA)-coated NPs causing the largest (85.8%) reduction (**Fig. 1c**). Moreover, TA-coated NPs exhibited ∼11.7-fold higher binding to RBCs compared to uncoated NPs, while the other coated NPs showed minimal enhancement (**Fig. 1d**). Notably, when taking the relative abundance of RBCs and WBCs into consideration, TA-coated NPs showed a >300-fold higher binding to RBCs over WBCs (**Fig. S1**). Compared to the other phenol compounds tested for NP coating, tannic acid contains a much higher number of galloyl units (8-10 per molecule). This multivalent nature enables simultaneous interactions with multiple sites on the RBC membrane which, in addition to the abundance of RBCs in whole blood, likely contributes to the higher RBC-binding efficiency of TA-coated NPs. Based on these results, we selected TA-coated NPs, referred to as i-Bind NPs hereafter, for subsequent studies.

Our transmission electron microscopy (TEM) analysis revealed a coating layer on the surface of i-Bind NPs, which was absent on uncoated PLGA NPs (**Fig. 1e**). Indicated by the cryo-TEM images (**Fig. 1f** and **Fig. S2**), i-Bind NPs showed a relatively rougher surface feature than uncoated PLGA NPs. To further characterize the coating, we performed Fourier transform infrared (FTIR) spectroscopy on both i-Bind and PLGA NPs (**Fig. 1g**). The FTIR spectrum of i-Bind NPs displayed characteristic peaks from both PLGA and TA, indicating the presence of TA on PLGA NPs. Notably, the broad OH-stretching peak between 3400 and 3050 cm^-1^, characteristic of TA, appeared in the i-Bind spectrum. Additionally, the C=O stretching peak at 1750 cm^-1^, characteristic of PLGA, was slightly shifted to 1749.57 cm^-1^ in i-Bind NPs, likely influenced by the TA-derived aromatic ester C=O stretch at 1698.14 cm^-1^. The aromatic ring stretching vibration of TA at 1604 cm^-1^ was also observed in i-Bind NPs, shifted slightly to 1612 cm^-1^. These data collectively suggest the successful coating of TA-metal complexes onto i-Bind NPs. Further characterization showed that i-Bind NPs had an average hydrodynamic diameter of 197.7 nm with a narrow size distribution (**Fig. 1h-i**). Moreover, i-Bind NPs are more negatively charged compared to PLGA NPs, likely due to ionization of the phenolic groups on TA (**Fig. 1j**). The colloidal stability of i-Bind NPs was evaluated over 15 days (**Fig. S3**). i-Bind NPs remained stable in dextrose saline (the injection vehicle) and serum protein containing buffers (PBS with 2% or 50% fetal bovine serum), with minimal changes in hydrodynamic size over time (**Fig. S3**). Notably, the larger NP size in protein containing buffers is likely due to protein binding to NP surfaces.

### i-Bind NPs *in situ* bind to RBCs

We evaluated the *in situ* binding efficiency of i-Bind NPs to RBCs under various conditions. We first assessed NP binding in purified RBC samples lacking serum proteins and other blood components. RBCs isolated from mouse whole blood were resuspended at 10% hematocrit and incubated with DiD-labeled NPs, which showed minimal free dye transfer to RBC membranes (**Fig. S4**), for 30 minutes under continuous rotation. NP binding to RBCs was then characterized by fluorescence imaging. i-Bind NPs, exhibited stronger binding to RBCs than uncoated PLGA NPs, as evidenced by the higher fluorescence intensity on RBC surface (**Fig. 2a-b**). To further quantify the efficiency of NP binding to RBCs, we fluorescently labeled RBCs with FITC-conjugated anti-Ter119 antibody and measured the binding of DiD-labeled NPs to RBCs by flow cytometry (**Fig. 2c**). The mean fluorescence intensity (MFI) of NPs on RBCs, an indicator of the number of NPs on RBCs, was significantly higher (1.5-fold) for i-Bind NPs compared to uncoated PLGA NPs (**Fig. 2d**), consistent with the fluorescence imaging results (**Fig. 2a-b**). We next examined the strength of the interaction between NPs and RBCs. After NP hitchhiking onto RBCs, samples were subjected to two cycles of centrifugation at 100 g for 5 minutes, equivalent to a shear stress of ∼ 3 Pa(*36*). The relative number of NPs remaining on RBCs was quantified by flow cytometry. While both PLGA and i-Bind NPs showed progressive detachment with washing, i-Bind NPs consistently retained a 4- to 18-fold higher number remaining on RBC surfaces compared to uncoated PLGA NPs after both washes (**Fig. 2d**), further indicating that i-Bind NPs had a greater binding affinity to RBC membrane. Furthermore, under a physiologically relevant hematocrit condition (40% hematocrit), i-Bind NPs also demonstrated a 5.67-fold higher binding to RBCs compared with PLGA NPs (**Fig. S5**).

**Fig. 2.**
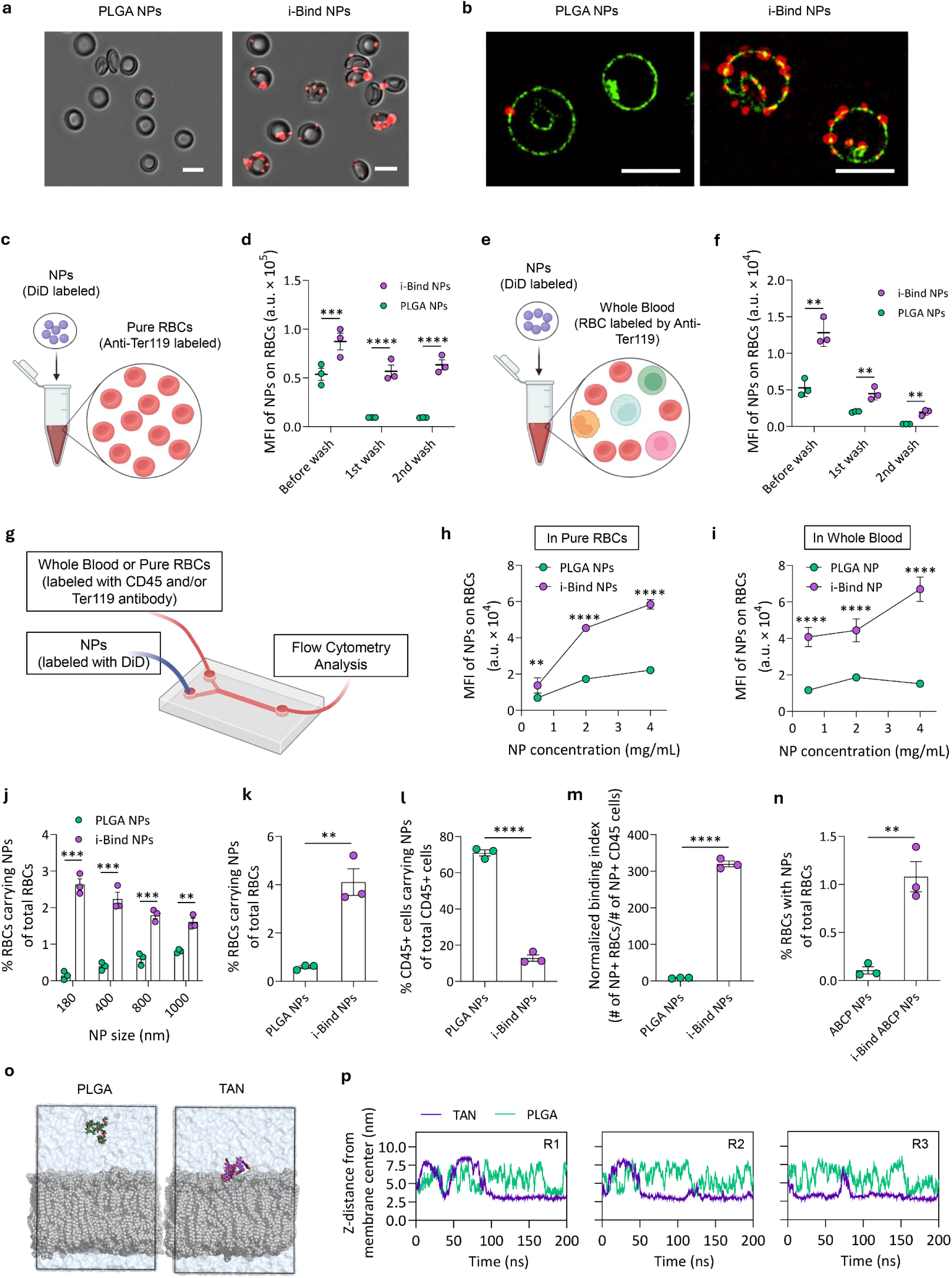
i-Bind NPs efficiently bind to RBCs in flowing whole blood conditions. **a,** Fluorescent microscopic images overlayed on brightfield images, showing binding of DiD-labeled PLGA or i-Bind NPs to RBCs at a RBC/NP incubation ratio of 400 to 1 in 10% pure RBC samples. Scale bars: 5 µm. **b,** High-resolution DeepSIM microscopic images showing the binding of DiD-labeled NPs (red color) onto RBCs in 10% pure RBCs. RBCs were pre-labeled with FITC-tagged anti-Ter119 antibody (green color). Scale bars: 5 µm. **c-d,** Characterization of NP binding to RBCs in pure RBCs. **c,** Schematic depicting the experimental setup for the evaluation of DiD-labeled NPs binding to RBCs in pure RBC samples. Created in BioRender. Zhao, Z. (2025) https://BioRender.com/q9rc2mv. **d,** Number of NPs (indicated by MFI) bound to RBCs after free mixing and following 2 wash-induced shearing in pure RBC samples (10% hematocrit). **e-f,** Characterization of NP binding to RBCs in mouse whole blood. **e,** Schematic showing the experimental design. Created in BioRender. Zhao, Z. (2025) https://BioRender.com/76otyy0. **f,** Number of NPs (indicated by MFI) bound to RBCs after free mixing and following 2 wash-induced shearing in mouse whole blood. **g-i,** Characterization of NP binding to RBCs in flowing pure RBC or whole blood conditions in a microfluidic device. **g,** Schematic showing the experimental setup. Created in BioRender. Zhao, Z. (2025) https://BioRender.com/rwpnw3u. **h-i,** Number of NPs (indicated by MFI) bound to RBCs in flowing 10% RBCs **(h)** or whole blood **(i)** at different feed NP concentrations. **j,** Binding of NPs with various sizes to RBCs (indicated by percentage of RBCs carrying NPs) in the microfluidic flowing whole blood condition. **k-m,** Characterization of NP binding to RBCs and CD45^+^ WBCs in the microfluidic flowing whole blood condition. Percentage of RBCs **(k)** and CD45^+^ WBCs **(l)** carrying NPs of total respective cells was shown. **m,** Relative binding index (ratio of NP^+^ RBC number to NP^+^ WBC number). **n,** Binding of uncoated or TA-coated ABCP polymeric NPs to RBCs in whole blood. **o-p,** Molecular dynamics simulations of the interaction of NP coating with RBC lipid membrane. **o,** Snapshots of the PLGA and tannic acid (TAN) interaction with RBC membranes in a simulation box (line). Ligands are depicted in green (PLGA) and purple (TAN) sticks, membrane atoms are in grey balls, water molecules are depicted as blue surface. The straight lines depict the edges of the simulation box. **p,** Time resolved distance evolution between the TAN and PLGA center of mass and the RBC membrane center of mass, projected to the z-axis. R1, R2, and R3 represent three independent replicates of simulations. Data in **(d, f, h-n)** are presented as mean ± SEM. For **(d, f, h-n**), statistical analysis was conducted by two-way ANOVA or two-tailed student’s t test: ** p < 0.01, *** p < 0.001, **** p < 0.0001.

We next assessed the RBC-binding efficiency of NPs in whole blood, which contains not only RBCs but also other blood cells and components (e.g., serum proteins) (**Fig. 2e**). Both i-Bind and PLGA NPs exhibited decreased binding to RBCs in whole blood compared to pure RBC samples, as indicated by decreased MFI of NPs on RBCs (**Fig. 2f**). This reduction is likely due to the formation of a protein corona, which interferes with NP binding kinetics and limits interactions with the RBC membrane(*37, 38*). Despite this, i-Bind NPs demonstrated significantly higher RBC-binding efficiency than uncoated PLGA NPs at the same dose (**Fig. 2f**). Flow cytometry analysis revealed that i-Bind NPs successfully hitchhiked onto 5.41% of RBCs compared to only 0.59% for PLGA NPs (**Fig. S6**) and showed a 2.4-fold higher MFI on RBCs (**Fig. 2f**). When binding strength was examined using centrifugal force as described previously, i-Bind NPs consistently retained a 2- to 6-fold higher number remaining on RBCs than uncoated PLGA NPs after both washes (**Fig. 2f**). Together, these results demonstrate that i-Bind NPs could maintain strong and effective binding to RBCs even in the complex environment of whole blood.

Next, we employed a bifurcated microfluidic chip to simulate blood flow and conducted a set of experiments to evaluate the capability of i-Bind NPs to bind to RBCs under dynamic whole-blood flow conditions (**Fig. 2g**). Using this setup, we evaluated how NP concentration and size influence their interaction with RBCs. i-Bind NPs showed 3- to 7-fold higher binding to RBCs across all the tested concentrations, as indicated by significantly higher MFI of NPs bound to RBCs in both 10% pure RBCs (**Fig. 2h**) and whole blood (**Fig. 2i**). Moreover, the interaction between i-Bind NPs and RBCs under flow conditions was observed to be dependent on the hydrodynamic size of the NPs. Across all four tested NP sizes (180-1000 nm), i-Bind NPs consistently showed significantly higher binding to RBCs than uncoated PLGA NPs in flowing whole blood conditions (**Fig. 2j**).

While tannic acid functionalization renders NP surface highly reactive and capable of interacting with a broad range of molecules, in this study, the preferential hitchhiking of i-Bind NPs on RBCs is likely driven by the high abundance of RBCs in whole blood, which increases the possibility of i-Bind NPs interacting with RBCs over interactions with other blood cell types. To test this, we investigated NP interaction with RBCs and CD45^+^ immune cells in whole blood. For this study, whole blood samples were first labeled with fluorescently labeled anti-Ter119 and anti-CD45 antibodies to distinguish RBCs and immune cells, respectively. DiD-labeled PLGA or i-Bind NPs were then incubated with mouse whole blood under microfluidic flow conditions as previously described. Compared to PLGA NPs, i-Bind NPs showed significantly increased binding to RBCs (**Fig. 2k**) and reduced binding to CD45^+^ WBCs (**Fig. 2l**). Notably, taking the relative blood cell abundance into consideration, i-Bind NPs showed a >300-fold higher binding to RBCs over WBCs (**Fig. 2m**), indicating i-Bind NPs’ preferential binding for RBCs over WBCs. Furthermore, to account for differences in cell abundance in whole blood, i-Bind NP interactions with RBCs and WBCs were evaluated under normalized cell number conditions (**Fig. S7a**). RBCs and WBCs were isolated by gradient centrifugation and incubated with i-Bind NPs separately for 3 minutes at the same cell and NP concentration. Under these conditions, i-Bind NPs exhibited significantly higher association with RBCs compared with WBCs (**Fig. S7b**). Next, to characterize the *in situ* RBC hitchhiking of i-Bind NPs *in vivo*, we quantified NP binding to circulating RBCs 2 minutes after intravenous administration. i-Bind NPs exhibited 18.2-fold higher binding to RBCs, further indicating their strong *in situ* RBC hitchhiking potential (**Fig. S8a-b**).

To elucidate the differential interactions of tannic acid-coated NPs (TAN) with RBCs and WBCs, we performed atomistic molecular dynamics (MD) simulations of tannic acid and reference PLGA coating interacting with asymmetric RBC and WBC membrane models. The membrane models were initially constructed in the absence of membrane-associated ligands to evaluate intrinsic lipid membrane properties. Both systems remained stable and maintained characteristic lipid bilayer behavior throughout the simulations. While both membranes exhibited highly negatively charged intracellular leaflets (**Fig. S9, Table S2**), the extracellular leaflet of WBCs was slightly negatively charged, whereas the RBC extracellular leaflet was neutral. The RBC membrane displayed a higher degree of lipid ordering (**Fig. S10**), consistent with its increased cholesterol and sphingomyelin content (**Table S2**). After the addition of the ligands to the simulation, comparison of NP coating binding behavior showed that PLGA remained primarily solvated in the aqueous phase, whereas TAN preferentially associated with RBC lipid membranes and remained membrane-bound for most of the simulation period (**Fig. 2o-p**). This enhanced membrane affinity was supported by hydrogen-bond analysis, which revealed negligible PLGA–membrane interactions but extensive hydrogen bonding between TAN and membrane lipids (**Table S3**). Structural analysis of TAN interactions identified the middle and outer gallic acid hydroxyl groups as the primary regions responsible for membrane binding, whereas the inner glucose and ester regions primarily interacted with water molecules (**Fig. S12, Table S4**). The hydroxyl groups formed preferential hydrogen bonds with lipid phosphate oxygens (**Fig. S13**), indicating that phosphate-containing lipids potentially serve as the major interaction sites for TAN.

We further examined the membrane-specific interactions of TAN with RBC and WBC models. Biased simulations yielded more negative binding free energy for TAN towards the WBC membrane than the RBC membrane (−4.4 vs. −2.9 kcal/mol) and deeper TAN insertion in the WBC membrane compared with the RBC membrane (**Fig. S11, Fig. S14**). This difference was attributed to the higher structural ordering and rigidity of the RBC membrane. Atomic-level analysis demonstrated that TAN binding was largely nonspecific with respect to lipid composition and was primarily mediated by interactions between gallic acid hydroxyl groups and lipid phosphate groups (**Fig. S13**). Accordingly, most TAN–membrane hydrogen bonds involved phosphate oxygens (**Table S5**), which are present in both glycerophospholipids enriched in WBC membranes and sphingomyelins abundant in RBC membranes. These results explain TAN’s broader association with cell membranes relative to PLGA and suggest that the experimentally observed preferential RBC binding is not driven by a specific lipid-binding mechanism, but rather by a general affinity for lipid surfaces, combined with the higher abundance of RBCs in circulation. The simplified membrane models used here lack membrane proteins, which are more abundant on WBCs, as well as other dynamic cellular components; therefore, additional physiological factors, including membrane organization, glycocalyx interactions, protein-mediated steric effects, and membrane stability, may influence NP accessibility and retention. These factors, together with increased total membrane lipids due to RBCs’ total larger membrane surface area, may contribute to the experimentally observed RBC preference.

To test the impact of metal ion selection of i-Bind on RBC binding, we prepared i-Bind NPs using a different metal ion, manganese. Notably, i-Bind NPs prepared from manganese-tannic acid complexation also exhibited consistent RBC binding, indicating that tannic acid-RBC interactions, rather than the specific metal ion, drive hitchhiking (**Fig. S15).** To further explore the applicability of the i-Bind approach to other polymeric NPs, we utilized an anionic bottle brush copolymer (ABCP)(*39, 40*), to synthesize NPs and evaluated their RBC-binding ability (**Fig. S16a-b**). Notably, unlike PLGA NPs, TA coating did not render ABCP NPs more negatively charged, suggesting that surface charge changes upon TA coating are not solely governed by phenolic group ionization but rather likely reflect the combined effects of TA ionization, polymer surface composition, molecular organization, hydration, and charge presentation. TA functionalization of the NPs resulted in significantly increased RBC binding in 10% purified RBCs (**Fig. S16c**), static whole blood (**Fig. 2n**), and microfluidic flow whole blood conditions (**Fig. S16d**). Moreover, we assessed hemolysis induced by i-Bind NPs at various concentrations and compared it with PLGA NPs. The hemolysis rate remained below 2% at concentrations up to 10 mg/mL, indicating the minimal adverse effects of i-Bind NPs to RBCs (**Fig. S17**).

### i-Bind NPs show preferential accumulation to the lungs

Previous studies have demonstrated that NPs hitchhiked on RBCs *ex vivo* are mechanically dislodged in the narrow lung capillaries upon intravenous injection and predominantly accumulate in the lungs, which is the first downstream capillary bed encountered after intravenous administration(*12, 17*). To evaluate the tissue-level distribution of i-Bind NPs, we conducted a series of biodistribution studies in both healthy mice and disease models. Five hours after intravenous injection in healthy mice, uncoated PLGA NPs primarily accumulated in the liver and spleen, whereas i-Bind NPs showed significantly higher deposition in the lungs and reduced accumulation in the liver (**Fig. 3a**). Specifically, i-Bind NPs exhibited a 3-fold decrease in liver accumulation and a 6-fold increase in lung accumulation (**Fig. 3b**), resulting in a 23-fold higher lung-to-liver accumulation ratio (**Fig. 3c**). We also directly compared i-Bind NPs with uncoated PLGA NPs hitchhiked onto RBCs *ex vivo* (*ex vivo* RBC-HH) administered at the same NP dose. *Ex vivo* RBC-HH NPs showed a ∼2-fold higher lung accumulation than i-Bind NPs (**Fig. S18**). This is likely because not all injected i-Bind NPs successfully hitchhike onto RBCs *in vivo*, leading to a lower efficiency compared to the *ex vivo* approach. However, the administrable NP dose of the *ex vivo* RBC-HH approach is limited by low NP hitchhiking efficiency (typically <20% of input) and the number of RBCs that can be injected. In contrast, i-Bind NPs leverage endogenous circulating RBCs *in situ* and thus allows for repeated dosing at higher NP doses.

**Fig. 3.**
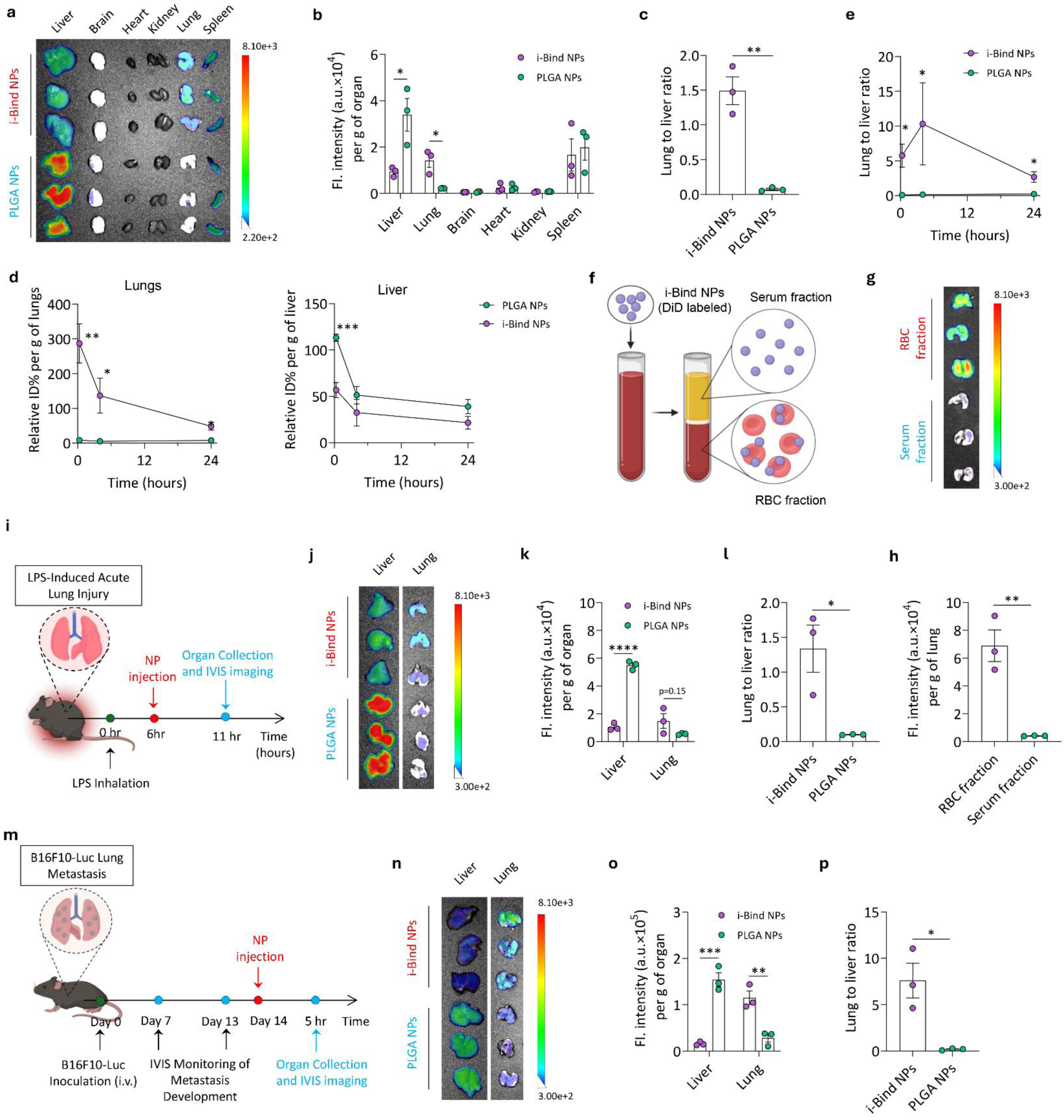
i-Bind NPs enable preferential delivery to lungs in healthy and disease conditions. **a-b,** Biodistribution of i-Bind NPs in healthy mice 5 hours after intravenous administration (n=3 biologically independent animals). **a,** Lago X images of major organs. **b,** Quantification of relative NP accumulation in different organs. **c,** Lung-to-liver ratio of NP accumulation. **d-e,** Time-course accumulation profiles of NPs in lungs and liver **(d)** and corresponding lung-to-liver accumulation ratio **(e)** after intravenous administration in healthy mice (n=4 biologically independent animals). **f-h,** Lung accumulation of i-Bind NPs is majorly contributed by RBC hitchhiking. **f,** Schematic showing experimental design to dissect the role of RBC hitchhiking versus intrinsic targeting of i-Bind NPs in lung accumulation. RBC fraction: i-Bind NPs hitchhiked on RBCs; serum fraction: i-Bind NPs not hitchhiked on RBCs and saturated with protein corona. Created in BioRender. Zhao, Z. (2025) https://BioRender.com/de22dig. **g,** Lago X image showing lung accumulation of i-Bind NPs 5 hours after intravenous injection of the serum or RBC portion. **h,** Quantification of NP accumulation in the lungs. **i-l,** Biodistribution of i-Bind NPs in an LPS-induced ALI model 5 hours after intravenous injection (n=3 biologically independent animals). **i,** Schematic showing experimental design. Created in BioRender. Zhao, Z. (2025) https://BioRender.com/ut0otug. **j,** Lago X images of liver and lungs of ALI mice. **k,** Quantification of NP accumulation in the liver and lungs. **l,** Lung-to-liver ratio of NP accumulation in the ALI model. **m-p,** Biodistribution of i-Bind NPs in a B16F10-Luc melanoma lung metastasis model 5 hours after intravenous injection (n=3 biologically independent animals). **m,** Schematic showing experimental design. Created in BioRender. Zhao, Z. (2025) https://BioRender.com/ut0otug. **n,** Lago X images of liver and lungs of mice with B16F10 lung metastasis. **o,** Quantification of NP accumulation in the liver and lungs. **p,** Lung to liver ratio of NP accumulation in the lung metastasis model. NPs in all the biodistribution studies were labeled with DiD. Data in **(b-e, h, k-l, o-p)** are presented as mean ± SEM. For **(b, c, d, e, h, k, l, o, p)**, statistical analysis was conducted by two-tailed student’s t test: * p < 0.05, ** p < 0.01, *** p < 0.001, **** p < 0.0001.

We further evaluated the pharmacokinetics of NPs in blood over 24 hours following intravenous administration. Both PLGA and i-Bind NPs showed rapid clearance from the circulation; notably, i-Bind NPs exhibited faster blood depletion than PLGA NPs (**Fig. S19a**). Within 2 min post-injection, circulating levels of i-Bind NPs decreased to ∼20% of the injected dose, whereas PLGA NPs remained above 80%. This accelerated decrease of i-Bind NPs in blood is likely due to their rapid association with RBCs, followed by detachment from RBCs and deposition in the lungs-the first downstream capillary bed after intravenous injection, reached within seconds to minutes. In a separate follow-up study, we measured the relative distribution of the remaining i-Bind NPs between serum and RBCs within the blood for 2, 10, or 30 minutes after intravenous administration. Despite their rapid overall decrease from blood, the majority (65-73%) of the i-Bind NPs remaining in circulation were associated with RBCs, further indicating their *in-situ* RBC hitchhiking capability (**Fig. S19b**). Next, we performed an extended biodistribution study in healthy mice to assess the tissue distribution kinetics of i-Bind NPs. At all tested time points, i-Bind NPs demonstrated a 7- to 30-fold increase in lung accumulation and an approximately 2-fold reduction in liver accumulation compared to uncoated PLGA NPs (**Fig. 3d** and **Fig. S20**), leading to a significantly higher lung-to-liver accumulation ratio (**Fig. 3e**). Notably, i-Bind NPs were rapidly redistributed from blood to the lungs within 20 mins and remained in the lungs for at least 24 hours, indicating their potential to enable prolonged lung-targeted drug delivery via systemic administration. To dissect the relative contribution of RBC hitchhiking versus intrinsic targeting to lung-specific accumulation of i-Bind NPs, we incubated i-Bind NPs with mouse whole blood for 10 mins to allow for RBC hitchhiking, separated the RBC and serum fractions carefully to minimize any potential fraction carry-over, and intravenously injected total each fraction separately into mice (**Fig. 3f**). The RBC fraction contained RBC-bound i-Bind NPs, while the serum fraction contained protein corona-saturated i-Bind NPs that did not associate with RBCs. Notably, the RBC fraction resulted in a 16.6-fold higher lung accumulation compared with the serum fraction (**Fig. 3g-h**). In a follow-up study, after separating the RBC and serum fractions, we intravenously injected the total serum fraction or part of the RBC fraction that contains equal dose of NPs. In this setting, the RBC fraction resulted in 4.87-fold higher lung accumulation of NPs compared with the serum fraction (**Fig. S21**). These data collectively indicate that RBC hitchhiking, rather than intrinsic targeting, is the dominant mechanism driving lung accumulation of i-Bind NPs. Moreover, we also tested the biodistribution of i-Bind NPs prepared by Mn-TA complexation (Mn-i-Bind NPs). Mn-i-Bind NPs showed >100-fold higher lung accumulation compared with PLGA NPs (**Fig. S22**), indicating the i-Bind approach can be applicable to metal ions beyond iron.

To evaluate the lung-targeting capability of i-Bind NPs in pathological conditions, we conducted biodistribution studies in two lung disease models: acute lung injury (ALI) and lung metastasis. In the lipopolysaccharide (LPS) inhalation-induced ALI model (**Fig. 3i**), i-Bind NPs showed a 5.2-fold reduction in liver accumulation and a 2.7-fold enhancement in lung accumulation compared to uncoated PLGA NPs (**Fig. 3j-k** and **Fig. S23**), resulting in a 13.3-fold higher lung-to-liver accumulation ratio (**Fig. 3l**). Similarly, in the B16F10 melanoma lung metastasis model (**Fig. 3m**), i-Bind NPs significantly enhanced lung accumulation and significantly reduced liver deposition relative to uncoated PLGA NPs (**Fig. 3n-p** and **Fig. S24**). Together, these findings collectively highlight the potential of i-Bind NPs for targeted drug delivery to the lungs under disease conditions.

Moreover, to assess the broader applicability of the i-Bind approach to other polymeric NPs, we examined the biodistribution of TA-coated ABCP NPs (i-ABCP NPs) in healthy mice. Intravenous administration of i-ABCP NPs resulted in significantly higher lung accumulation than uncoated ABCP NPs, yielding a 30-fold increase in the lung-to-liver ratio (**Fig. S25a-c**). These findings demonstrate that the i-Bind approach is a versatile strategy for delivering polymeric NPs to lungs via *in situ* RBC hitchhiking.

### i-Bind NPs preferentially target lung immune cell subsets

To further investigate the cellular-level distribution of i-Bind NPs in the lungs, we performed flow cytometry analysis on dissociated lung cells. Lungs were collected 5 hours after intravenous NP administration, processed into single-cell suspensions, and stained with fluorescently labeled antibodies to identify various cell populations. Uptake of i-Bind NPs by each cell type was then quantified by flow cytometry. Both i-Bind and uncoated PLGA NPs were predominantly taken up by CD45^+^ immune cells compared to endothelial and epithelial cells (**Fig. 4a**). Compared to uncoated PLGA NPs, i-Bind NPs showed enhanced uptake across all three cell categories, with the highest enrichment observed in CD45^+^ immune cells (4.1-fold increase) (**Fig. 4a**). Further analysis of the CD45^+^ immune cell population revealed that i-Bind NPs were distributed among multiple innate immune cell types and exhibited significantly increased uptake in CD11b^+^ type 2 conventional dendritic cells (cDC2s), CD103^+^ type 1 DCs (cDC1s), and neutrophils, with 1.6- to 4.9-fold enhancement relative to uncoated PLGA NPs (**Fig. 4b**). Among these, the highest uptake of i-Bind NPs was seen in CD11b^+^ cDC2s. This preferential accumulation may be attributed to the role of cDC2s in immune surveillance, as they actively sample the lung environment for antigens(*41, 42*).

**Fig. 4.**
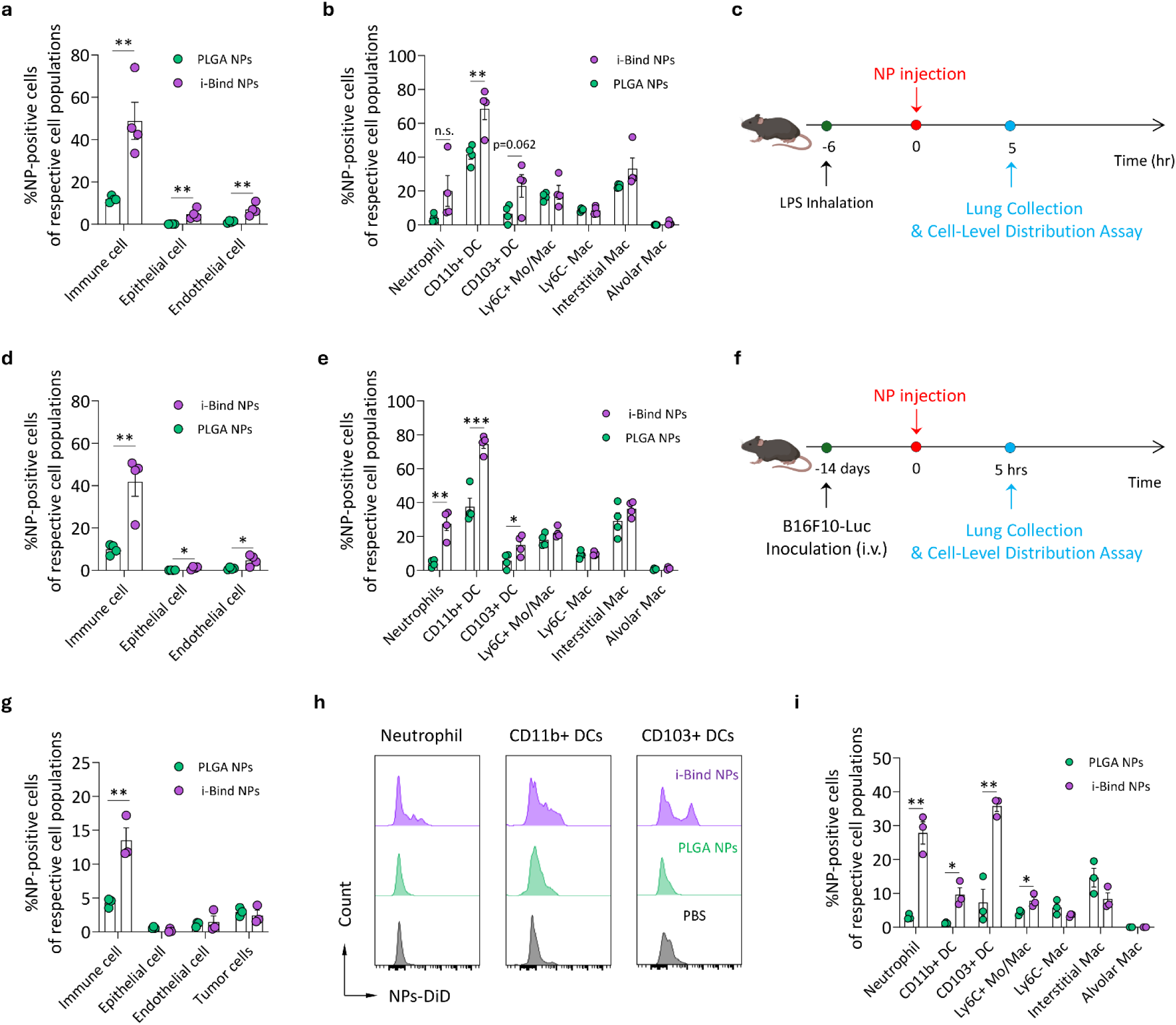
i-Bind NPs predominantly target resident immune cells in the lungs. **a-b,** Cell-level distribution of NPs in different cell types in the lungs of healthy mice 5 hours after intravenous administration (n=4 biologically independent animals). **a,** Percentage of different cells associated with NPs in the lungs. **b,** Percentage of various CD45^+^ immune cells associated with NPs in the lungs. **c-e,** Distribution of NPs in different cell types in the lungs of ALI mice 5 hours after intravenous administration (n=4 biologically independent animals). **c,** Schematic depicting the study design. Created in BioRender. Zhao, Z. (2025) https://BioRender.com/ut0otug. **d,** Percentage of different cells associated with NPs in the ALI lungs. **e,** Percentage of various CD45^+^ immune cells associated with NPs in the ALI lungs. **f-i,** Distribution of NPs in different cell types in the lungs of mice with lung metastasis 5 hours after intravenous administration (n=3 biologically independent animals). **f,** Schematic illustration for the study design. Created in BioRender. Zhao, Z. (2025) https://BioRender.com/ut0otug. **g,** Percentage of different cells associated with NPs in the lungs with metastases. **h,** Representative flow cytometry histograms showing the accumulation of NPs in neutrophils, CD11b^+^ DCs, and CD103^+^ DCs. **i,** Percentage of various CD45^+^ immune cells associated with NPs in the lungs with metastases. In (**b, e, i**), DC, Mo and Mac refer to dendritic cell, monocyte, and macrophage, respectively. Data in **(a-b, d-e, g, i)** are presented as mean ± SEM. For **(a-b, d-e, g, i)**, statistical analysis was conducted by two-tailed student’s t test: * p < 0.05; ** p < 0.01; *** p < 0.001; n.s.: no statistically significant difference.

As the immune landscape differs substantially between healthy and diseased lungs, we further investigated the cellular-level distribution of i-Bind NPs under ALI and lung metastasis conditions. In the murine ALI model (**Fig. 4c**), similar to the observations in healthy lungs, i-Bind NPs were predominantly associated with CD45^+^ immune cells, although they also exhibited increased uptake by lung endothelial and epithelial cells (**Fig. 4d**). Further analysis revealed that, compared to PLGA NPs, i-Bind NPs showed significantly higher accumulation in neutrophils in addition to CD11b^+^ cDC2s (**Fig. 4e**). This neutrophil accumulation is likely driven by the increased neutrophil infiltration during acute lung injury, triggered by chemokine signaling from damaged epithelial cells and resident macrophages. As first responders in ALI, neutrophils play a central role in pathogen clearance via phagocytosis(*43, 44*), which may underline their enhanced uptake of i-Bind NPs. In the B16F10 melanoma lung metastasis model (**Fig. 4f**), i-Bind NPs again preferentially accumulated in CD45^+^ immune cells (**Fig. 4g**), consistent with observations in both healthy and ALI lungs. In addition to CD11b^+^ cDC2s, i-Bind NPs also showed enhanced accumulation in CD103^+^ cDC1s, neutrophils, and Ly6C^low^ monocytes/macrophages, with the highest enrichment seen in CD103^+^ cDC1s (**Fig. 4h-i**). cDC1s are a specialized subset of conventional DCs that play a critical role in anti-tumor immunity by cross-presenting antigens to cytotoxic CD8 T cells(*45, 46*). Targeting and modulating cDC1s in the lungs with metastases enabled by i-Bind NPs may provide a new approach for developing more effective immunotherapies for lung metastasis treatment. Together, our results comparing healthy and disease conditions suggest that i-Bind NPs enhance delivery to the lung microenvironment, where they are subsequently taken up by immune cell subsets that are enriched and/or activated under specific pathological conditions. While previous studies have demonstrated NP loading onto RBCs *ex vivo* for targeting the lungs and associated immune cells(*17, 34*), our *in-situ* approach extends this approach by demonstrating disease-specific localization. We show that the distribution of immune cells and, consequently the i-Bind NP association vary with disease state, indicating that our i-Bind strategy can potentially achieve targeted delivery that responds to pathological conditions rather than relying solely on baseline lung targeting.

### i-Bind NPs carrying a STING agonist potently inhibit lung metastasis

Next, we evaluated the therapeutic potential of i-Bind NPs in the treatment of lung diseases. Leveraging their lung- and immune cell-targeting capabilities, we assessed the anti-metastatic efficacy of diABZI-loaded i-Bind NPs (referred to as i-Bind diABZI NPs) in a B16F10 lung metastasis model (**Fig. 5a**). diABZI is a potent STING agonist currently under investigation in early-stage clinical trials for solid tumors. It functions primarily by activating innate immune cells, dendritic cells and macrophages in particular, within the tumor microenvironment to elicit anti-tumor immune responses(*47, 48*). diABZI was encapsulated into PLGA NPs with an encapsulation efficiency of 8.45%, corresponding to a loading capacity of 16.9 µg diABZI per mg of NPs. The physicochemical properties of the NPs were characterized (**Fig. S26a**), and the diABZI i-Bind NPs remained stable over 8 days (**Fig. S26b**). *In vitro* drug release analysis demonstrated sustained diABZI release from NPs, with approximately 21% cumulative release after 72 h from PLGA NPs. diABZI i-Bind NPs exhibited a significantly slower release profile with 1.5% cumulative release at 72 h, possibly because the tannic acid coating provides an additional barrier that limits drug diffusion and reduces premature diABZI leakage from the PLGA NP matrix.

**Fig. 5.**
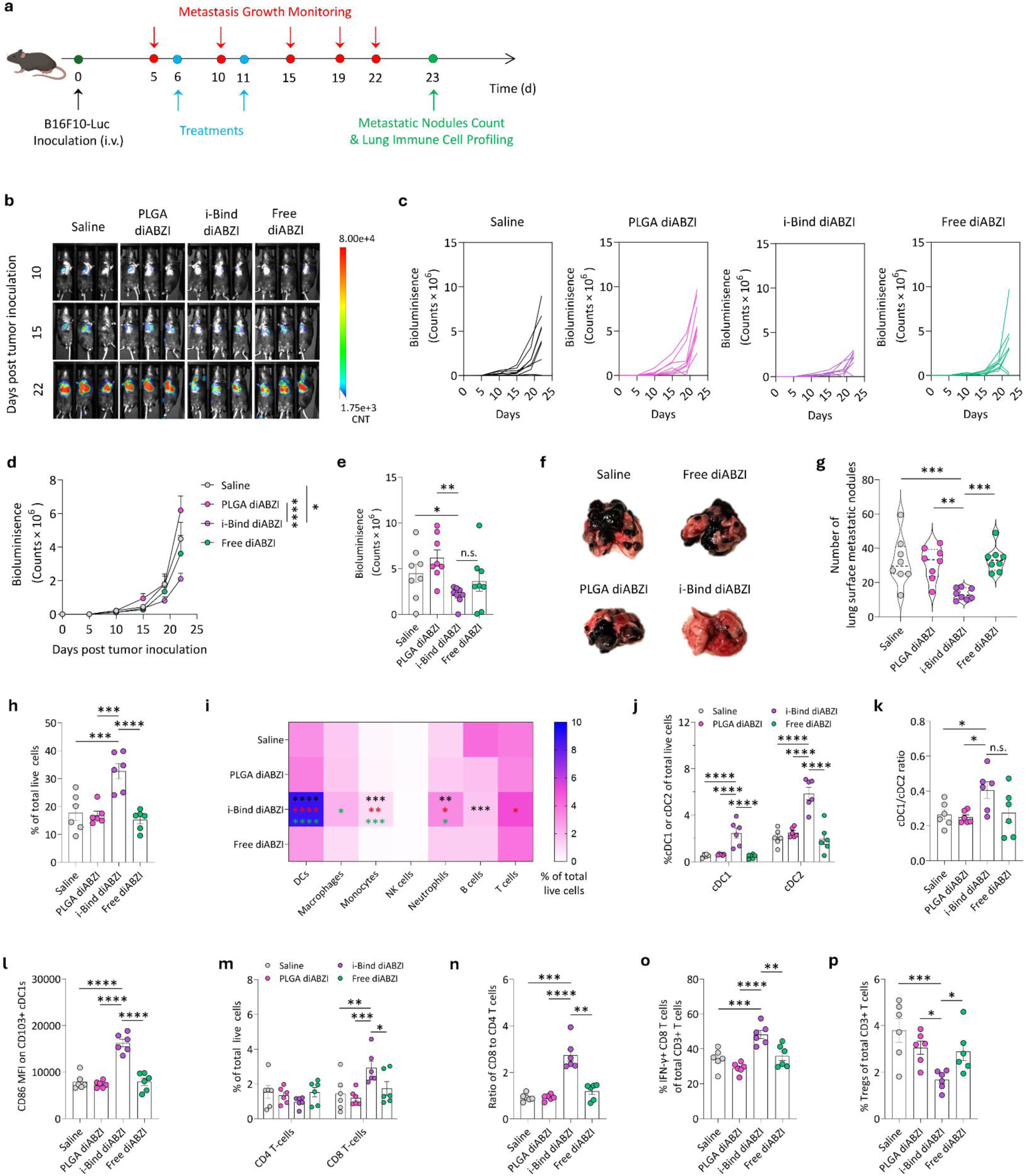
i-Bind NPs encapsulating a STING agonist reprogram the lung immune microenvironment and inhibit lung metastasis progression. **a-g,** Therapeutic efficacy of i-Bind NPs loaded with diABZI and other control formulations (n=8 biologically independent animals). **a,** Schematic illustration of the treatment schedule. Created in BioRender. Zhao, Z. (2025) https://BioRender.com/ut0otug. **b,** Representative bioluminescence images of lung metastasis progression at different time points. **c,** Lung metastasis progression of individual mouse indicated by bioluminescence intensity. **d,** Overall lung metastasis progression curve. **e,** Lung metastasis tumor burden on day 22 indicated by bioluminescence intensity. **f,** Representative images of lungs following different treatments on day 23. **g,** Number of surface metastatic nodules on the lungs on day 23. **h-p,** Changes of the lung immune cell profiles following different treatments on day 23 (n=6 biologically independent animals). **h-i,** Number of total CD45^+^ immune cells **(h)** and various immune cell subtypes **(i)** in the lungs. **j,** Number of CD103^+^ cDC1s and CD11b^+^ cDC2s in the lungs. **k,** Relative ratio of cDC1s to cDC2s. **l,** Expression level of DC activation marker CD86 on cDC1s (indicated by Mean fluorescence intensity, MFI). **m,** Number of CD4 and CD8 T cells in the lungs. **n,** Relative ratio of CD8 to CD4 T cells in the lungs. **o,** Relative number of IFN-γ-expressing CD8 T cells of total CD3 T cells in the lungs. **p,** Relative number of Tregs of total CD3 T cells in the lungs. Data in **(d, e, g-h, j-p)** are presented as mean ± SEM. For (**d**), statistical analysis was conducted by two-way ANOVA: * p < 0.05, **** p < 0.0001. For **(e, g-i, j-p)**, statistical analysis was conducted by one-way ANOVA: * p < 0.05, ** p < 0.01, *** p < 0.001, **** p < 0.0001; n.s. indicates no statistically significant difference between indicated groups. For (**i**), black, pink, and green asterisks indicate comparisons between the i-Bind diABZI group with saline, PLGA diABZI, and free diABZI groups, respectively.

In our therapeutic efficacy study, mice received two doses of treatment and were monitored for disease progression over 23 days (**Fig. 5a**). Treatment efficacy was evaluated based on lung bioluminescence intensity derived from luciferase-expressing tumor cells. As shown in the imaging and quantification data (**Fig. 5b-d** and **Fig. S27**), i-Bind diABZI NPs significantly inhibited the progression of lung metastases, while free diABZI and PLGA diABZI NPs showed minimal therapeutic benefits. By day 22, i-Bind diABZI NPs reduced the average tumor burden by 2.2-, 2.9-, and 1.7-fold compared to saline, PLGA diABZI NPs, and free diABZI, respectively (**Fig. 5e**). Further analysis on day 23 confirmed the efficacy of i-Bind diABZI NPs, leading to a 2.5-fold reduction in the number of metastatic nodules on the lung surface (**Fig. 5f-g** and **Fig. S28a**) and a 1.5-fold decrease in lung weight compared to the saline treatment (**Fig. S28b**). Collectively, these results demonstrate that i-Bind NPs significantly enhanced the therapeutic efficacy of diABZI in treating lung metastases, likely due to their ability to effectively deliver the therapeutic cargo, diABZI, to lungs and immune cells. Notably, blank i-Bind NPs without diABZI loading did not induce obvious activation of dendritic cells (CD103⁺ cDC1s) *in vitro*, suggesting that the NP carrier itself contributes minimally to immune activation and that the therapeutic efficacy is primarily driven by diABZI delivery (**Fig. S29).**

We then performed immune cell profiling of metastatic lung tissues on day 23 post-tumor inoculation to elucidate the immunological mechanisms underlying the therapeutic efficacy of i-Bind diABZI NPs. While free diABZI and PLGA diABZI NP treatments did not show obvious effect, i-Bind diABZI NP treatment led to a substantial increase in CD45^+^ immune cell infiltration into metastatic lungs, showing a 2.1-fold enhancement compared to saline-treated controls (**Fig. 5h**). Further analysis revealed that i-Bind diABZI NPs also significantly altered the composition of immune cells in metastatic lungs (**Fig. 5i**). In saline-treated lungs, B cells were the predominant immune cell population. However, i-Bind diABZI NP treatment shifted this immune cell landscape toward a DC-and T-cell dominant profile. Notably, i-Bind diABZI NP treatment resulted in DCs accounting for 50% of the total immune cells in the lungs while also significantly increasing the infiltration of macrophages, monocytes, neutrophils, and T-cells compared to saline and other control treatments (**Fig. 5i**).

Activation of the STING pathway plays a crucial role in innate immune response, resulting in the activation of DCs by promoting their maturation as well as upregulating the expression of antigen-presenting molecules (MHCI and MHCII), co-stimulatory molecules (e.g., CD80 and CD86), and migratory chemokine receptor CCR7(*49*). This enables DCs to process antigens and present them to T-cells in the lymph node for eliciting an antigen-specific adaptive immune response(*50, 51*). We further analyzed the activation status and subtype of DCs in the metastatic lungs following different treatments. The lung-targeted delivery of diABZI using i-Bind NPs resulted in the recruitment of 4.7-fold more cDC1s and 2.8-fold more cDC2s to the metastatic lungs compared to the saline treatment, while free diABZI and PLGA diABZI NPs did not show obvious effect (**Fig. 5j**). Notably, i-Bind diABZI NPs significantly increased the relative ratio of cDC1s to cDC2s (**Fig. 5k**). CD103^+^ cDC1s have been recognized as a crucial immune cell population for the transport of antigens to the lymph node, priming of cytotoxic CD8 T cells, and the recruitment of T cells to the tumor microenvironment(*46*). Typically, this distinctive DC subtype has limited presence due to the tumor’s immunosuppressive environment. Interestingly, i-Bind diABZI NPs were the only treatment to recruit significantly higher numbers of CD103^+^ cDC1s to the lungs (**Fig. 5j**), likely due to their capability to enhance diABZI delivery to the lungs. Furthermore, i-Bind diABZI NPs resulted in significantly stronger CD86 activation of cDC1s (**Fig. 5l**). This observation highlights i-Bind diABZI NPs as an effective approach for STING agonist-induced activation of DCs in the lung metastasis model.

We also analyzed the T cell profiles in the metastatic lungs to assess the efficacy of i-Bind NP-enhanced diABZI delivery in modulating adaptive immunity. We observed notable differences in the CD4 and CD8 T cell populations of the lungs treated by i-Bind diABZI NPs. While free diABZI and PLGA diABZI NPs showed minimal effect on the lung infiltration of T cells, i-Bind diABZI NPs led to a 40% reduction in CD4 T cell numbers but caused a 2.0-fold enhanced infiltration of CD8 T cells compared to the saline treatment (**Fig. 5m**). This resulted in a 2.3-fold increase in CD8/CD4 T cell ratio compared to the saline group (**Fig. 5n**). These data are in line with our DC data showing that i-Bind diABZI NPs enhanced the infiltration and activation of DCs, particularly CD103^+^ cDC1s, to the lungs which likely led to more efficient priming and expansion of CD8 T cells and infiltration of CD8 T cells to the lungs. Additionally, further analysis revealed that i-Bind diABZI NPs also significantly increased the number of effector (IFN-γ^+^) CD8 T cells which are effective in tumor cell killing and significantly reduced the number of immunosuppressive regulatory T cells (Tregs) in the metastatic lungs (**Fig. 5o-p**). Collectively, our lung immune cell profiling data indicate that i-Bind NP-mediated enhancement of diABZI delivery to innate immune cells in metastatic lungs is associated with immune activation and remodeling of the tumor microenvironment toward a more immunologically “hot” state, along with suppression of lung metastasis progression. Notably, while many studies have shown the ability of STING agonists to induce proinflammatory responses and increase the recruitment of effector T-cells, their application in treating metastasis is limited by their ineffective accumulation to the metastatic tissues(*52, 53*). The i-Bind strategy offers a promising approach for the effective delivery of diABZI and STING agonist-mediated modulation of lung-resident immune cells, leading to increased T cell infiltration and activation that is not achieved with free diABZI or diABZI-loaded PLGA NPs. Collectively, our data suggest that i-Bind NPs, by enabling targeted delivery of therapeutic agents to innate immune cells, can effectively reprogram the diseased lung microenvironment and offer a promising strategy for treating lung disorders associated with immune dysfunction.

To assess the safety and biocompatibility of blank or diABZI-loaded i-Bind NPs, we performed blood chemistry and hematological analyses on healthy mice 7 days post intravenous administration (**Fig. S30** and **Fig. S31**). Overall, administration of i-Bind NPs, either blank or diABZI-loaded, did not result in significant changes in the evaluated complete blood count and blood chemistry parameters compared with the PBS treatment, suggesting the overall safety of the i-Bind platform. Notably, diABZI-loaded i-Bind NPs slightly increased monocyte count and decreased lymphocyte and platelet count, most likely due to the effect of the payload diABZI, as blank i-Bind NPs did not lead to these trends.

## Discussion

In this study, we reported an *in situ* RBC-binding nanoparticle (i-Bind NP) that spontaneously associates with RBCs in flowing whole blood and enables efficient lung targeting. This platform leverages tannic acid surface functionalization of polymeric NPs to form multiple non-covalent interactions with RBC membranes via polyphenolic groups. RBC hitchhiking is an emerging strategy in which NPs anchored to RBC surface *ex vivo* exploit the natural circulation of RBCs to extend systemic circulation time and achieve targeted organ accumulation. Although RBCs are readily available and their reinfusion is well-established clinically, the technical complexity associated with *ex vivo* attachment of therapeutic cargos to RBC membranes remains a major barrier to clinical translation. Subtle alterations introduced during e*x vivo* manipulation may lead to suboptimal delivery outcomes and unwanted adverse reactions. Moreover, *ex vivo* RBC hitchhiking is not easily scalable or adaptable to repeated dosing. Our i-Bind approach provides a simple and efficient solution to overcome these challenges, enabling *in situ* RBC hitchhiking of NPs under physiological blood flow to achieve lung-preferential targeting. While i-Bind is less effective than the *ex vivo* approach in lung delivery at equal NP doses, it allows flexible dose adjustment, personalization, and repeat dosing without repeated RBC manipulation. While the i-Bind approach may still share some similar or even more pronounced limitations as the *ex vivo* approach, such as immunogenicity due to RBC modification and off-target modification of other blood cells (e.g., leukocytes, platelets), it eliminates the deleterious effects of *ex vivo* RBC manipulation during the manufacturing process and the immunogenic risks of donor RBCs bearing foreign antigens. As a result, *in situ* RBC hitchhiking via i-Bind transforms RBC-mediated delivery from a cell-based intervention into a true injectable drug platform.

Earlier *ex vivo* RBC hitchhiking work with polymeric nanoparticles, such as PLGA NPs, suggested their attachment to RBCs through non-covalent interactions; however, these particles could not efficiently hitchhike in whole blood and tend to detach even under low shear stress, limiting their *in vivo* performance(*15, 54*). In contrast, our *in vitro* binding assays showed that tannic acid-coated i-Bind NPs bind to RBCs in significantly greater numbers than uncoated PLGA NPs, and that the tannic acid coating resulted in increased RBC binding and reduced WBC association compared to other polyphenol coatings. These findings are consistent with our molecular dynamics simulation results, which revealed that the tannic acid coating of i-Bind NPs exhibits broadly enhanced affinity towards RBC lipid membranes via hydrogen-bonding interactions between the gallic acid hydroxyl groups of tannic acid and lipid phosphate groups. Notably, tannic acid contains 8-10 galloyl groups per molecule, while the other polyphenols evaluated in this study contain only 1-2. This higher galloyl content and hydroxyl density lead to greater multivalent hydrogen-bonding capacity, facilitating faster and more robust RBC binding compared to the other polyphenols tested. The other polyphenols with weaker membrane-binding capacity associate less stably with the RBC surface, rendering them more available for internalization by WBCs via endocytosis or phagocytosis, likely contributing to the elevated WBC interaction observed with other coatings. Under flow conditions, i-Bind NPs demonstrated a significant increase in RBC hitchhiking, attributable to the adhesive properties of tannic acid coating. Mechanistically, the i-Bind strategy does not rely on specific receptor-ligand recognition but instead on broadly enhanced affinity to RBC membranes. Our simulation results indicate that the tannic acid coating has broad, non-differentiating affinity for both RBC and WBC lipid membranes, suggesting that the RBC-preferential binding observed in our wet-lab experiments is unlikely to stem from an intrinsically stronger tannic acid affinity for RBC versus WBC lipid membranes. Instead, this preference most likely arises from the generally enhanced lipid membrane affinity of tannic acid combined with key differences in abundance, biophysical characteristics, and NP association mechanisms between RBCs and WBCs. First, RBCs are far more abundant than WBCs in whole blood (∼1000:1 ratio) and collectively present a much greater total membrane surface area, thereby increasing the likelihood of NP-RBC interactions. RBC membranes also exhibit greater structural order and rigidity than WBC membranes. Together, these factors favor NP retention on RBCs once initial association occurs. Second, WBCs possess more membrane-associated proteins and engage in dynamic cellular processes (e.g., endocytosis/phagocytosis) that can limit NP accessibility. NP association with RBCs is rate-limited by NP-membrane binding, whereas association with WBCs is rate-limited by endocytosis/phagocytosis, a process considerably slower than membrane binding. This difference in overall association kinetics favors preferential hitchhiking of tannic acid-coated NPs onto RBCs. These two factors collectively enable efficient *in situ* RBC hitchhiking in whole blood. Although independent of specific receptor recognition, i-Bind enables reproducible association in whole blood, which is advantageous for physiologically relevant and scalable systemic transport. Consistent with these findings, biodistribution studies revealed significantly higher lung accumulation of i-Bind NPs in both healthy and diseased mice. Additionally, our extended biodistribution data demonstrate that within 20 min, i-Bind NPs achieve peak accumulation in the lungs, gradually decreasing over 24 h, indicating spontaneous RBC binding and efficient lung targeting without concerns of prolonged systemic circulation and toxicity. Given that the pulmonary vasculature receives the entire cardiac output and comprises ∼30% of total endothelial cells(*55, 56*), lung deposition is often attributed to RBC-endothelium contact. Surprisingly, while i-Bind NPs did accumulate in endothelial cells, a greater proportion localized to CD45^+^ immune cells. In naïve mice, uptake was observed in cDC2 dendritic cells; in acute lung injury (ALI), neutrophil uptake predominated; and in a lung metastasis model, cDC1 dendritic cells were the primary targets. These patterns likely reflect the prevailing immune cell composition in each condition, suggesting that i-Bind NPs could be adapted to preferentially deliver immunomodulatory agents to distinct immune populations. This distribution is driven by a two-step mechanism. First, i-Bind NPs preferentially and quickly bind to RBCs after intravenous administration. When NP-carrying RBCs squeeze through the narrow lung capillaries, which is the first downstream capillary bed, i-Bind NPs get mechanically dislodged and deposit within the pulmonary capillary bed. Second, the subsequent NP accumulation in immune cells is likely driven by local immune surveillance and active sampling of the lung microenvironment by resident and recruited immune cell populations. Therefore, rather than directly targeting specific immune subsets, i-Bind NPs may enhance delivery to disease-relevant immune populations by increasing NP availability within the affected tissue environment. While previous *ex vivo* RBC hitchhiking work focused on broad lung delivery, we demonstrate that i-Bind NP distribution reflects disease-induced immune cell dynamics, providing an opportunity for disease-tailored cargo customization.

In the lung metastasis model, delivering only two doses of diABZI via i-Bind NPs at a subtherapeutic dose (2 mg/kg) significantly reduced metastatic nodule formation and delayed disease progression compared to free drug. This effect was likely driven by enhanced drug accumulation in lung immune cells, particularly cDC1s, which, in turn, promoted greater recruitment and activation of cytotoxic T cells than either free drug or PLGA NPs. Furthermore, i-Bind NP treated mice exhibited reduced regulatory T cell (Treg) levels, indicating a shift toward a less immunosuppressive microenvironment. The enhanced activation of both innate and adaptive immune responses observed with i-Bind diABZI NPs compared with free diABZI highlights their potential for immunomodulatory therapy. Notably, in the current study, we only assessed i-Bind NP-mediated diABZI delivery efficacy through lung immune cell profile changes and did not directly measure STING pathway activation. We acknowledge that direct molecular readouts, such as IFN-β, CXCL10, pTBK1, or pIRF3, would provide additional mechanistic evidence of pathway activation, and this will be investigated in future studies.

Overall, our results demonstrate that i-Bind NPs can spontaneously bind to RBCs after systemic administration, enabling preferential deposition to immune cells in the lungs. This reduces off-target exposure and maintains therapeutic presence in the lungs. Notably, previous studies have shown that altering the injection site in RBC hitchhiking can direct NPs to tissues immediately downstream of the injection vessel, such as the brain and kidneys(*17*). Accordingly, although the present study focuses on lung targeting, i-Bind could potentially serve as a generalized targeting strategy for NPs in other tissues by simply varying the injection location. Taken together, the i-Bind strategy offers a simple, versatile, and adaptable approach to enhance organ-preferential targeting of polymeric NPs via *in situ* RBC hitchhiking, with the potential to tailor immune cell targeting and reprogramming across diverse pathological conditions. It is important to note that other elegant NP surface coating strategies for *in situ* RBC hitchhiking, such as ionic liquid-based coatings, have also been reported(*18, 25*). This approach employs tunable interfacial interactions to promote NP association with RBC membranes *in vivo*. The relative effectiveness of i-Bind versus these other strategies remains to be explored in future studies. Notably, in this work, we have focused on testing i-Bind in mouse models; the applicability of i-Bind in larger animal models and humans remains to be tested. Additionally, future studies should validate the i-Bind approach in spontaneous rather than experimental lung metastasis models, as well as other lung disease models. Furthermore, while our *ex vivo* and *in vitro* data demonstrate preferential binding of i-Bind NPs to RBCs over WBCs, the dynamics of these interactions within the *in vivo* circulation remain to be fully elucidated. Future studies will focus on evaluating RBC–NP interactions *in vivo* to better understand and optimize the behavior of i-Bind NPs under physiological conditions. Moreover, while our serum/RBC fraction transfer experiments demonstrated that RBC hitchhiking, rather than intrinsic NP targeting, is the major driver of i-Bind NP lung accumulation, tannic acid coating likely alters multiple NP physicochemical properties such as size, surface charge, and roughness. The role of these changes in i-Bind NP RBC binding and lung accumulation remains to be fully elucidated in future studies.

## Materials and Methods

### Materials

Poly(D,L-lactic-co-glycolide) (PLGA, Resomer RG653H), polyvinyl alcohol (PVA), tannic acid (TA), iron(III) chloride hexahydrate, and lipopolysaccharide (LPS, from *Escherichia coli* O111:BA) were purchased from Sigma-Aldrich (St. Louis, MO). STING agonist diABZI (Compound 3) was purchased from Selleck Chemicals LLC (Houston, TX). Gallic acid, 4-morpholinepropanesulfonic acid (MOPs), caffeic acid, epigallocatechin gallate, and catechin hydrate were purchased from Thermo Fisher Scientific (Waltham, MA). 1,1’-Dioctadecyl-3,3,3’,3’-Tetramethylindodicarbocyanine (DiD), 3,3’-Dioctadecyloxacarbocyanine Perchlorate (DiO), 1,1’-Dioctadecyl-3,3,3’,3’-Tetramethylindocarbocyanine Perchlorate (DiI) were purchased from Thermo Fisher Scientific (Waltham, MA). High Activity General Tissue Enzymatic Digestion Kit (DHGT-5004) was purchased from RWD Life Science (Dover, DE). Penicillin/streptomycin was obtained from Cytiva (Marlborough, MA). High glucose-Dulbecco’s modified eagles’ medium (DMEM) and heat-inactivated fetal bovine serum (FBS) were purchased from Corning (Corning, NY). Antibodies including Zombie NIR (#423105), Spark Blue554 anti-CD45 (#103184), AF700 anti-CD45 (#157615), PE anti-Ly6G (#164503), FITC anti-Ly6C (#128006), BV650 anti-CD11c (#117339), PerCP-Fire806 anti-CD11b (#101294), BV605 anti-MHCII (#107639), PE/Dazzle594 anti-CD64 (#139319), PE/CY5 anti-CD24 (#101811), PE/Fire700 anti-CD3 (#100271), Spark UV387 anti-CD4 (#100491), AF700 anti-CD19 (#115527), PE/CY7 anti-CD49b (#108921), APC-Fire810 anti-CD25 (#102076), AF647 anti-FoxP3 (#126408), PerCp/CY5.5 anti-CD206 (#141716), APC-Fire750 IFN-γ (#505860), Pacific blue anti-CD8 (#100725), PE anti-CD31 (#106203), FITC anti-CD326 (#118207), PE/CY7 anti-CD170 (Singlec-F) (#155528), BV605 anti-CD11b (#101237), APC/CY7 anti-CD11c (#117323), PerCp/CY5.5 anti-Ly6C (#128011), Pacific Blue anti-Ly6G (#127611), Spark UV387 anti-MHCII (#107669), PE anti-TER119 (#116207), FITC anti-TER119 (#116206), and PE/CY5 anti-CD24 (#101811) were all purchased from BioLegend (San Diego, CA).

### Preparation of PLGA NPs

PLGA NPs were prepared using a double emulsion method(*16, 57*). In brief, 20 mg of PLGA (Resomer RG653H, Mw 24,000-38,000, 65:35 L:G ratio) was dissolved in 1 mL of dichloromethane (DCM) as the organic phase to which 150 µL of phosphate-buffered saline (PBS) was added. This was followed by sonication in a water bath (Bransonic Ultrasonic Corporation, CT, USA) for 1 min to form the primary emulsion. The primary emulsion was added dropwise to 11 mL of 1.5% polyvinyl alcohol (PVA) (w/v) under magnetic stirring at 1200 rpm and subsequently probe sonicated (Fisherbrand Model FB505, USA) at 50% magnitude for 40 seconds with a 20-second break. The organic phase was removed by evaporation under stirring in a fume hood for 12 hours. Fluorescently labeled PLGA NPs were prepared using a similar method by adding 50 µg DiD in the organic phase prior to the formation of the primary emulsion. To obtain PLGA NPs of various sizes, the PVA concentration, PLGA mass, and sonication settings were varied accordingly: 180 nm NPs (0.5% PVA; probe sonication: 40 seconds on, 20 seconds pause, 50% sonication magnitude); 300-400 nm NP (0.1% PVA; probe sonication: 40 seconds on, 20 seconds pause, 50% sonication magnitude); 500-600 nm NP (2% PVA; homogenizer: level 1, 1 minute) and 1000 nm NP (2% PVA; homogenizer: level 3, 5 minutes).

### Preparation of ABCP NPs

ABCP, an anionic bottlebrush copolymer, was produced following a previously published method(*39, 40*). Briefly, it was synthesized using a combination of ring-opening and controlled radical polymerizations from a backbone of glycidyl methacrylate (GMA) and 2-(2-bromoisobutyryloxy)ethyl methacrylate (BIEM). The hydrophobic block side-chains consisted of poly(D,L-lactide) (LA), whereas the hydrophilic block side-chains consisted of a copolymer of poly(methacrylic acid) (MAA) and poly(2-methacryloyloxyethyl phosphorylcholine) (MPC) in an equimolar ratio. The chemical structure of ABCP [poly((GMA111-g-LA14)-b-(BIEM105-g-(MPC12-co-MAA31))] and its molecular weight (Mn=1.15 MDa) were determined by 1H NMR, and its polydispersity (Đ=1.35) was measured by size exclusion chromatography.

To make ABCP NPs, ABCP polymer was dissolved in an organic phase composed of DMF/MeOH 1:1 v/v with DiD dye, and polymer to DiD dye mass ratio is consistent with previous PLGA NPs. The organic phase and aqueous phase (deionized water) have a volume ratio of 1:3 and were mixed through a home-made dual-inlet microfluidic device at flow rates of 10 mL/min and 30 mL/min, respectively, to generate 2 mg/mL NPs. The collected NPs were purified by dialyzing against a 100-fold volume of deionized (DI) water in a 3.5-5 kD dialysis membrane bag. DI water was changed every 2 hours for 2 times to remove the organic solvent.

### Preparation of i-Bind NPs

To prepare i-Bind NPs through polyphenol coating, PLGA NPs were washed in DI water three times and then resuspended to 1 mg/mL in DI water. To the suspension, 5 µL of phenolic compounds (e.g., 24 mM tannic acid) in DI water was added and vortexed for 10 seconds, followed by the addition of 5 µL iron chloride solution (24 mM) under bath sonication for 1 minute. Then, 50 µL of 4-morpholinepropanesulfonic acid (MOPs) was added and vortexed for 10 seconds, and the resulting polyphenol-coated NPs (i-Bind NPs) were washed three times with DI water and resuspended in dextrose saline (9:1 v/v mixture of 5% Dextrose and 0.9% saline). Coated ABCP NPs were prepared using a similar method. For manganese-i-Bind NPs, iron chloride was replaced with 10 µL of 24 mM manganese chloride solution under bath sonication for 1 minute. This was followed with adding 50 µL MOPs and 500 µL PBS and the mixture vortexed for 30 minutes. The resulting Mn-i-Bind NPs were washed three times with DI water and resuspended in dextrose saline.

### Characterization of NPs

To characterize the physicochemical properties of NPs, NPs were resuspended in ultrapure water at 1 mg/mL. The hydrodynamic size, polydispersity index, and surface charge of NPs were determined by a dynamic light scattering method, using a Malvern Nano-ZS Zetasizer (Malvern Instruments). The measurements were performed in MilliQ water at 25 ℃. Fourier-transform infrared (FTIR) spectroscopy was performed on lyophilized NP samples using a Nicolet^TM^ 6700 FTIR in the range of 500-4000 cm^-1^, under a setting of 16 scans at a resolution step of 4 cm^-1^. Transmission electron microscopy (TEM) using a JEM-1011 transmission electron microscope (JEOL) at 80 kV were carried out to characterize the morphology of NPs. A suspension of NPs in DI water was deposited on a Cu grid (150 mesh) coated with an ultrathin amorphous carbon film, and the drop was allowed to dry before analysis. For stability testing, NPs were resuspended and stored in dextrose saline or PBS supplemented with FBS (2% or 50%). The hydrodynamic diameter (Z-average) and polydispersity index (PDI) of the NPs were measured over a 15-day period.

### Characterization of NP binding to RBCs *in vitro*

All *in vitro* NP binding experiments were performed on freshly collected whole blood samples or purified RBCs from C57BL/6 mice. To obtain purified RBCs, the serum and buffy coat layer of whole blood were removed by centrifugation at 1000 g for 10 minutes at 4 °C. The resulting RBCs were washed with PBS three times before resuspending to 10% or 40% hematocrit in PBS or dextrose saline (9:1 v/v mixture of 5% dextrose and saline). NP suspension was added to the purified RBCs or whole blood in a self-standing screwcap tube and mixed thoroughly using a pipette. The samples were then rotated for 30 minutes on a tube revolver at 14 rpm. Subsequently, unbound NPs were removed by centrifuging samples at 100 g for 5 minutes at 4 ℃, and the samples were subsequently washed with PBS. To allow for quantification, the RBCs were fluorescently labeled with anti-Ter119 antibody (FITC, Biolegend, clone TER-119), and CD45+ WBCs were stained with anti-CD45 antibody (Alexa Fluor 700, Biolegend, clone QA17A26). Fluorescent microscopy and super-resolution imaging were used for qualitative determination of the NP binding efficiency to RBCs, while flow cytometry (Cytek AURORA) was used to quantify the binding of NPs on RBCs. To evaluate binding under flowing whole blood conditions, RBCs and WBCs were fluorescently labeled with an anti-Ter119 antibody (FITC, BioLegend, clone TER-119) and an anti-CD45 antibody (Alexa Fluor 700, BioLegend, clone QA17A26), respectively, and NPs were labeled with DiD. A 60° (30° + 30°) symmetrical bifurcated microfluidic chip (Synvivo, 100 μm parent, 50 μm daughter widths, 100 μm depth) was utilized, and blood and NP flow were maintained at 80 μL/min and 20 μL/min, respectively, using a syringe pump (New Era Pump Systems Inc., NY, USA). After each run, 20 μL of the sample was collected, diluted to 200 μL using dextrose saline buffer, and NP binding was quantified using flow cytometry. To evaluate NP binding affinity toward WBC and RBC populations under normalized cell number conditions, whole blood was collected and separated using density centrifugation with a Percoll gradient. The WBC and RBC fractions were carefully isolated and washed twice with PBS. The isolated cell populations were then stained with anti-CD45 antibody to identify WBCs and DiO membrane dye to label RBCs prior to incubation with fluorescently labeled NPs. Equal numbers of RBCs and WBCs were counted and incubated with same concentration of NPs separately for 3 minutes and NP binding to cells were analyzed by flow cytometry.

### Molecular dynamics simulations

The details concerning MD simulation can be found in Supporting Information.

### Super-resolution imaging

Fluorescence super-resolution imaging of RBC-NP binding was performed using a multispot structured illumination system (DeepSIM X-LIGHT, CrestOptics) coupled to an inverted microscope (Nikon Instruments, Eclipse Ti2E). A 60x/1.42 oil objective was used to acquire images, with a Kinetix sCMOS camera (serial number: A23H723002). Images were captured using the NIS-Elements software (version 5.42.04). Live-cell volumetric movies were captured using the “Deep” DeepSIM mode with 637 nm and 476 nm laser illumination and a step size of 0.5 µm.

### Hemolysis assay

Freshly isolated RBCs were resuspended to 5% hematocrit and incubated with different concentrations of NPs at room temperature for 2 hours. RBCs incubated in PBS and DI water were used as negative control and positive control, respectively. The samples were spun down at 500 g for 3 minutes to remove RBCs and 20,000 g for 15 minutes to remove NPs, and the supernatant was collected. Hemoglobin levels in the supernatant were detected by measuring absorbance at 540 nm using a SpectraMax® M3 Multi-Mode Microplate Reader (Molecular Devices). The hemolysis rate was calculated using the formula:

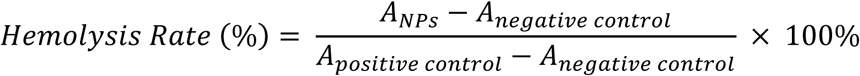

### Cell line and animals

*In vivo* studies were performed using C57BL/6 mice (6-7 weeks of age, female) purchased from Jackson Laboratory (ME, USA). All animal experiments were performed according to protocols (21–098, 24–085) approved by the Animal Care Committee (ACC) of the University of Illinois Chicago. The B16F10 melanoma cells expressing luciferase (B16F10-Luc) were generously gifted by Dr. Quanyin Hu from the University of Wisconsin-Madison and cultured in high-glucose Dulbecco’s modified Eagle’s medium (DMEM) supplemented with 10% fetal bovine serum (FBS), 1% penicillin-streptomycin, and 1 μg/mL puromycin. Cells were incubated in a humidified atmosphere with 5% CO_2_ at 37 ℃.

### *In vivo* biodistribution studies

To investigate the biodistribution of NPs, 100 μL of DiD-labeled NPs (5 mg/mL in dextrose saline) was intravenously administered via tail vein injection to C57BL/6 mice. The mice were sacrificed 5 hours post NP administration, and the NP accumulation within major organs (liver, lungs, spleen, kidneys, brain, and heart) was imaged and semi-quantified by Lago X (Spectral Instruments). To evaluate the circulation kinetics of NPs, mice were intravenously administered with DiD-labeled NPs via tail vein injection, and blood samples were collected at pre-determined time points post-injection. The DiD fluorescence intensity of blood samples was measured in black 96-well plates using SpectraMax® M3 Multi-Mode Microplate Reader (Molecular Devices). The zero-time point was determined by measuring the fluorescence signal of a known amount, equivalent to the injected dose, of DiD-labeled NPs diluted in blood collected from untreated control mice. The fluorescence intensity at the zero-time point was defined as 100%, and subsequent measurements were normalized and expressed as a percentage of the zero-time point signal. To determine the distribution of i-Bind NPs between blood fractions over time, blood samples collected at each time point were centrifuged at 300 g for 5 min. The RBC and serum fractions were subsequently collected, and the amount of NPs in each fraction was determined by measuring DiD fluorescence intensity. For an extended biodistribution study, Lago X imaging of major organs was performed at 20 minutes, 4 hours, and 24 hours post-injection, and the percent injected dose per gram of organ (%ID/g) was semi-quantitatively calculated based on the Lago X imaging data. %ID in one specific organ was calculated by dividing the fluorescence value of the interested organ by the total fluorescence value from all major organs of the same mouse. *In vivo* biodistribution of Mn-i-Bind NPs was evaluated similarly.

To characterize the association of NPs with RBCs *in vivo*, circulating RBCs were labeled before NP administration. To achieve *in vivo* RBC labelling, mice received 100 µL of filtered anti-mouse CD16/32 antibody (BioLegend, #101301) by tail-vein injection to reduce nonspecific Fc receptor-mediated antibody binding. After 15 min, 10 µg of FITC-anti-TER119 antibody (BioLegend, #116205; clone TER-119) and 10 µg of Pacific Blue-rat IgG2b, κ isotype-control antibody (BioLegend, #400627) were administered intravenously via tail vein. Following a 1-hour period, DiD-labeled NPs were administered by tail-vein injection. Blood was collected by 2 min after NP administration and immediately processed for flow cytometric analysis. RBCs were identified as TER119-positive/isotype-negative cells, and NP association with RBCs was quantified based on the DiD fluorescence signal.

To compare the biodistribution of *ex vivo* red blood cell–hitchhiked PLGA NPs (*ex vivo* RBC-HH) and i-Bind NPs, RBCs were isolated from freshly collected whole blood. Whole blood was centrifuged at 1,000 g for 10 min at 4 °C, after which the plasma and buffy coat layers were removed. The isolated RBCs were washed three times with PBS and resuspended in PBS to a hematocrit of 10%. To prepare *ex vivo* RBC-HH, 150 µL of DiD-labeled NP suspension (18.5 mg/mL) was added to 150 µL of the RBC suspension in a self-standing screwcap tube. The mixture was thoroughly mixed by pipetting and rotated for 40 min at 14 rpm using a tube revolver. Unbound NPs were subsequently removed by centrifugation at 100 g for 5 min at 4 °C. The RBCs were then washed twice with PBS and resuspended in 135 µL of PBS. NP loading onto the RBCs was determined by measuring DiD fluorescence intensity. For the biodistribution study, *ex vivo* RBC-HH or i-Bind NPs, each containing an equivalent NP dose of 10 µg, were intravenously administered to C57BL/6 mice via tail-vein injection. At 5 h post-injection, the mice were euthanized, and major organs were collected. NP accumulation in the excised organs was imaged and semi-quantified using a Lago X imaging system (Spectral Instruments).

To further characterize the RBC-hitchhiking potential of i-Bind NPs *in vivo*, whole blood was collected and incubated with DiD-labeled i-Bind NPs for 10 minutes. Subsequently, the blood was centrifuged at 300 g for 5 minutes, and the RBC and serum fractions were carefully collected to minimize any potential fraction carry-over. The resulting fractions were administered intravenously via the tail vein to mice representative of the RBC- and Serum groups. The mice were euthanized 5 hours post-NP administration, and the NP accumulation within major organs (liver, lungs, spleen, kidneys, brain, and heart) was imaged and semi-quantified by Lago X (Spectral Instruments). To further assess the RBC-hitchhiking potential under normalized NP conditions, following incubation, NP content in the RBC and serum fractions was quantified by fluorescence measurement using a SpectraMax® M3 Multi-Mode Microplate Reader (Molecular Devices). RBC volume with an NP amount equivalent to that in the serum fraction was used for the subsequent biodistribution study.

For evaluation of NP biodistribution in an acute lung injury (ALI) model, mice were placed in a restrainer and received aerosolized lipopolysaccharide (LPS) at a concentration of 5 mg/mL in PBS for 30 minutes. Subsequently, NPs were intravenously administered 5 hours after LPS inhalation. Major organs were collected 5 hours after NP injection and imaged using Lago X. For evaluation of biodistribution in a murine melanoma lung metastasis model, the lung metastasis model was established by intravenous injection of 0.1 million B16F10-Luc cells to C57BL/6 mice. On day 14, the establishment of lung metastases was confirmed by bioluminescence imaging using Lago X before NP administration. Major organs were collected 5 hours after NP injection and imaged using Lago X.

To evaluate the cellular level distribution of the NPs within the lung tissue, the lungs were collected 5 hours after DiD-labeled NP administration and processed to single cell suspensions using a Mouse High Activity General Tissue Enzymatic Digestion Kit (RWD, USA). Single cells were labeled with antibodies including AF700 anti-CD45 (Biolegend, #157615), PE anti-CD31 (Biolegend, #106203), FITC anti-CD326 (Biolegend, #118207), PE/CY7 anti-CD170 (Singlec-F) (Biolegend, #155528), BV605 anti-CD11b (Biolegend, #101237), APC/CY7 anti-CD11c (Biolegend, #117323), PerCp/CY5.5 anti-Ly6C (Biolegend, #128011), Pacific Blue anti-Ly6G (Biolegend, #127611), Spark UV387 anti-MHCII (Biolegend, #107669), PE/CY5 CD24 (Biolegend, #101811) and analyzed by flow cytometry (Cytek AURORA) to quantify NP distribution within different types of cells.

### Therapeutic efficacy and immune cell profiling studies

The therapeutic efficacy of diABZI loaded i-Bind NPs was studied in a B16F10-Luc melanoma lung metastasis model. diABZI-loaded i-Bind NPs were prepared using a similar double emulsion method as previously described, but the organic phase was switched to chloroform. Specifically, 2 mg of diABZI (Selleck Chemicals) was dissolved in 1 mL of chloroform as the organic phase and heated to 40 ℃ for 5 minutes to get a clear solution before making the primary emulsion. NPs were characterized as described previously, and the loading efficiency of diABZI in NPs was analyzed using ultra-high performance liquid chromatography (UHPLC) equipped with a C18 column, using a 5-95% gradient mobile phase consisting of acetonitrile (0.1% formic acid) in water (0.1% formic acid). To evaluate the *in vitro* drug release, diABZI-loaded NPs (1 mg) were washed and resuspended in 1 mL PBS containing 1% FBS, then rotated continuously at room temperature for 72 h. At predetermined time points, samples were centrifuged to pellet the NPs, and 200 µL of supernatant was collected and replaced with fresh PBS containing 1% FBS. Release samples were mixed 1:1 with an internal-standard solution containing Compound 2 and analyzed by LC–MS. Cumulative diABZI release was calculated after correcting for the sampled and replenished volumes. Chromatographic separation was performed on a Waters Xbridge C8 column (50 × 2.1 mm, 3.5 µm) at 40 °C and a flow rate of 0.3 mL/min. The mobile phases consisted of water with 0.1% formic acid (solvent A) and acetonitrile with 0.1% formic acid (solvent B). The separation gradient was from 30%-95% solvent B over 4 min. Quantitative analysis was done on a Shimadzu 8060 triple quadrupole mass spectrometer using positive ion electrospray with selected ion monitoring detection.

For the therapeutic efficacy study, 0.1 million B16F10-Luc cells were intravenously administered to female C57BL/6 mice (6-8 weeks of age, n=8 per group) to induce lung metastases. Mice were imaged using Lago X and randomized into 4 groups (saline, PLGA-diABZI, i-Bind diABZI, and free diABZI) based on the bioluminescence intensity of lung metastases on day 5. Two doses of diABZI formulations (equivalent to 2 mg/kg diABZI) were intravenously administered on days 6 and 11 post tumor inoculation, and the tumor burden was monitored on days 5, 10, 15, 19, and 22 by bioluminescence imaging using Lago X. On day 23, mice were euthanized, and major organs were collected. Tumor burden in the lungs was further evaluated by weighing the lungs and counting metastatic nodules on the lung surface.

To evaluate immune cell infiltration into lungs following different treatments, lungs were collected from mice on day 23. For the immune profiling study, six out of eight mice per group were randomly selected and analyzed, with two mice per group reserved for histological evaluation. Collected lung samples were further processed to single cells using a Mouse High Activity General Tissue Enzymatic Digestion Kit (RWD, USA), stained with an antibody panel consisting of Zombie NIR (Biolegend, #423105), Spark Blue 554 anti-CD45 (Biolegend, #103184), PE anti-Ly6G (Biolegend, #164503), FITC anti-Ly6C (Biolegend, #128006), BV650 anti-CD11c (Biolegend, #117339), PerCP-Fire806 anti-CD11b (Biolegend, #101294), BV605 anti-MHCII (Biolegend, #107639), PE Dazzle 594 anti-CD64 (Biolegend, #139319), PE/CY5 anti-CD24 (Biolegend, #101811), PE/Fire700 anti-CD3 (Biolegend, #100271), Spark UV387 anti-CD4 (Biolegend, #100491), AF700 anti-CD19 (Biolegend, #115527), PE/CY7 anti-CD49b (Biolegend, #108921), APC-Fire810 anti-CD25 (Biolegend, #102076), AF647 anti-FoxP3 (Biolegend, #126408), PerCp/CY5.5 anti-CD206 (Biolegend, #141716), APC-Fire750 anti-IFNγ (Biolegend, #505860), and Pacific Blue anti-CD8 (Biolegend, #100725), and analyzed by flow cytometry (Cytek AURORA). Data were analyzed using FlowJo 10.

### *In vitro* immune activation study

CD103^+^ cDC1s were cultured using a previously reported method(*57*). To evaluate DC activation by blank and diABZI-loaded NPs *in vitro*, CD103^+^ DCs were seeded in a 12-well plate (0.5 × 10^6^ cells per well) and cultured overnight. Subsequently, 200 µg/mL NPs or 1 µg/mL lipopolysaccharide were added to the cells, followed by further incubation for 12 h. The cells were then harvested, stained with antibodies against activation markers (MHCII, CD80, CD86), and analyzed by flow cytometry (Cytek AURORA).

### Hematological and blood chemistry study

For hematological and serum chemistry analyses, mice were administered 0.5 mg of i-Bind NPs, 0.5 mg of i-Bind diABZI NPs, or saline via tail-vein injection. At 7 days post-administration, blood was collected by retro-orbital sampling into heparinized tubes for hematological analysis or serum-separation tubes for serum chemistry analysis. Serum samples were allowed to coagulate for 20 min and then centrifuged at 400 g for 15 min. The serum was collected and transferred to clean sample tubes. Whole-blood and serum samples were shipped on ice to the IDEXX BioAnalytics laboratory (North Grafton, MA) for preclinical testing using short toxicity panel (#60513) and complete blood count panel (#61330).

### Statistical analysis

All experiments were repeated at least twice. Statistical analyses were performed using GraphPad Prism 10, and all data were presented as mean ± s.e.m. The level of statistical significance was determined using Student’s t-test, One-way ANOVA, or Two-way ANOVA. The levels of significance were depicted with asterisks as follows: p < 0.05 *; p < 0.01 **; p < 0.001 ***, p < 0.0001 ****. All the flow cytometry analyses were carried out using FlowJo 10 software.

## Acknowledgments

We thank Dr. Balaji Baskaran Ganesh from the flow cytometry core at UIC for helping with flow cytometry analysis. PC, MP, and MO also acknowledge VSB-Technical University of Ostrava, IT4Innovations National Supercomputing Center, Czech Republic for granting access to the LUMI supercomputer owned by the EuroHPC Joint Undertaking, hosted by CSC (Finland) and the LUMI consortium with the support of the Ministry of Education, Youth and Sports of the Czech Republic through e-INFRA CZ (grant 90254). COST Action CA21101 is also acknowledged.

## Funding

National Institutes of Health/National Institute of General Medicine R35GM150507 (ZZ)

National Institutes of Health/National Cancer Institute R21CA291723 (ZZ)

National Institutes of Health/National Heart, Lung, and Blood Institute R21HL168650 (ZZ)

Chancellor’s Translational Research Initiative Award from the Office of Technology Management at the University of Illinois Chicago (ZZ)

National Institutes of Health/National Institute of General Medicine R35GM146786 (YSH)

National Science Foundation (Grant No. 2003789) (MH)

Czech Science Foundation (Grant no. 25-18467S) (MO)

European Union REFRESH-Research Excellence for Region Sustainability and High-tech Industries project (No. CZ.10.03.01/00/22_003/0000048)- ERDF/ESF project TECHSCALE (No. CZ.02.01.01/00/22_008/0004587) (MO)

## Author contributions

Conceptualization: ZZ, EMU

Methodology: ZZ, EMU

Investigation: EMU, EZ, MMN, SH, HG, BF, CC, JL, LZ, PC, MP, DSN, MO

Visualization: EMU, ZZ

Supervision: ZZ

Writing—original draft: EMU, ZZ

Writing—review & editing: EMU, ZZ, EZ, MMN, SH, HG, BF, CC, LZ, JL, MH, PC, MP, DSN, MO, YL, YSH, ZP

## Data and materials availability

The main data supporting the results of this study are available in the main text or the supplementary materials. Data related to molecular dynamics simulations can be found at https://doi.org/10.5281/zenodo.20622319. Additional information is available for research purposes upon reasonable request from the corresponding author.

