## Supplementary information for "Lung-Selective Immune Reprogramming via *In Situ* Red Blood Cell Hitchhiking Nanoparticles"

**This PDF file includes:**

Supplementary Methods
Figs. S1 to S34
Table S1 to S5

**Other Supplementary Materials for this manuscript include the following:**

**Supplementary Methods**

**Molecular dynamic simulation studies**

The red blood cell (RBC) and white blood cell (WBC) lipid membrane systems were assembled using the CHARMM-GUI platform(*1, 2*) with composition shown in **Table S2**. The membranes were built using previously described Membrane Builder(*3-5*) and the ligands were parametrized using the Ligand Reader(*6*). The forcefield used was CHARMM36m(*7*) for lipids and CGenFF(*8*) for ligands. For water, TIP3P (Transferable Intermolecular Potential 3P)(*9*) with the default $Na^{+}$ and $Cl^{-}$ ion parameters and the WYF cation-pi reparameterization(*10*) were also used. The composition of the RBC membranes was based on the experimental composition previously determined(*11*), while the composition of WBC membranes was determined based on reported measurements(*12-16*). Following the area-per-lipid (APL) process previously described(*17*), symmetric membranes of the extracellular/intracellular leaflets of the studied membranes were built with the desired composition, subject to a simulation run, and then the asymmetric membranes were assembled using area per lipid calculated from the symmetric membranes. All membranes were subjected to the CHARMM-GUI equilibration protocol, with stepwise restraint release and a simulation step increase from 0.001 ps to 0.002 ps. The production run was for 200 ns at 303.15 K with V-rescale temperature coupling(*18*), semi-isotropic C-rescale pressure coupling set to 1 bar(*19*) and LINCS(*20*) constraints applied to all hydrogen atom bonds.

The ligands were placed into the center of a box of water 3 nm high, which was placed on top of the final snapshot structure of the equilibrated asymmetric membranes, using the GROMACS gmx solvate and gmx editconf tools. The complex systems with ligands were minimized with the steepest descent algorithm for 25,000 steps and then subject of 200 ns of production simulation with the same settings as above. Each system was run in three replicates. The last 50 ns of the simulation run (sampled every 100 ps) was used for numerical analyses. The free energy profiles were calculated by umbrella simulations. Starting from the TAN's deepest embedded structure from unbiased simulations, we pulled TAN towards the membrane center using a pull rate of 0.1 nm/ns and an umbrella potential of 10,000 kJ/mol/nm. Umbrella simulation starting structures were extracted with TAN positions ranging from the middle of the membrane up to fully solvated in water (5 nm), spaced by 0.1 nm. For each structure, a biased simulation with an umbrella potential of 2,000 kJ/mol/nm was performed for 50 ns per simulation window. The final free energy profile was extracted with the weighted histogram analysis method(*21*), discarding the initial 20 ns for equilibration. The simulations were run in GROMACS 2023.1(*22-24*) and analyzed in MDAnalysis 2.9.0(*25, 26*), LiPyphilic 0.12.1(*27*), and gorder 1.2.0 (*28*). Their dependencies are Freud(*29*) and hydrogen bond analysis methodology previously described(*30*).

**Supplementary Figures**

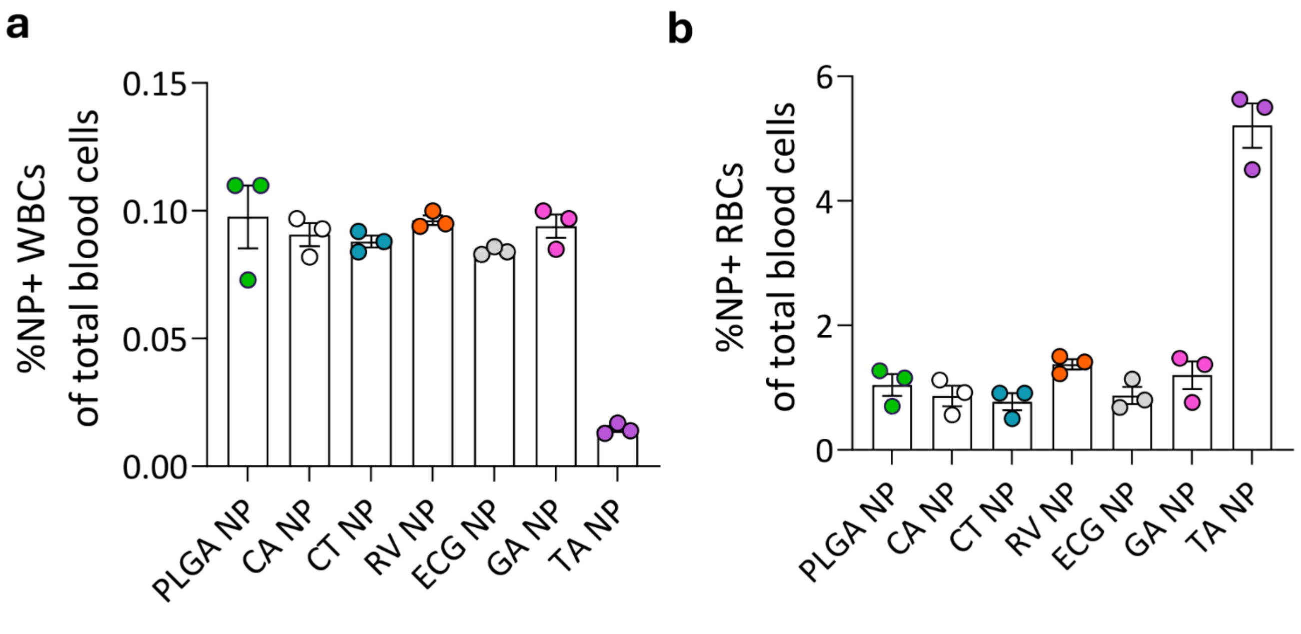


**Fig. S1. Binding of polyphenol-coated NPs to CD45⁺ WBCs and RBCs in whole blood when accounting for the relative abundance of RBCs and WBCs in whole blood. a,** Percentage of NP+ WBCs of total blood cells. **b,** Percentage of NP+ WBCs of total blood cells. CA: caffeic acid; CT: catechin; RV: resveratrol; ECG: epigallocatechin gallate; GA: gallic acid; TA: tannic acid. Data are presented as mean ± SEM.

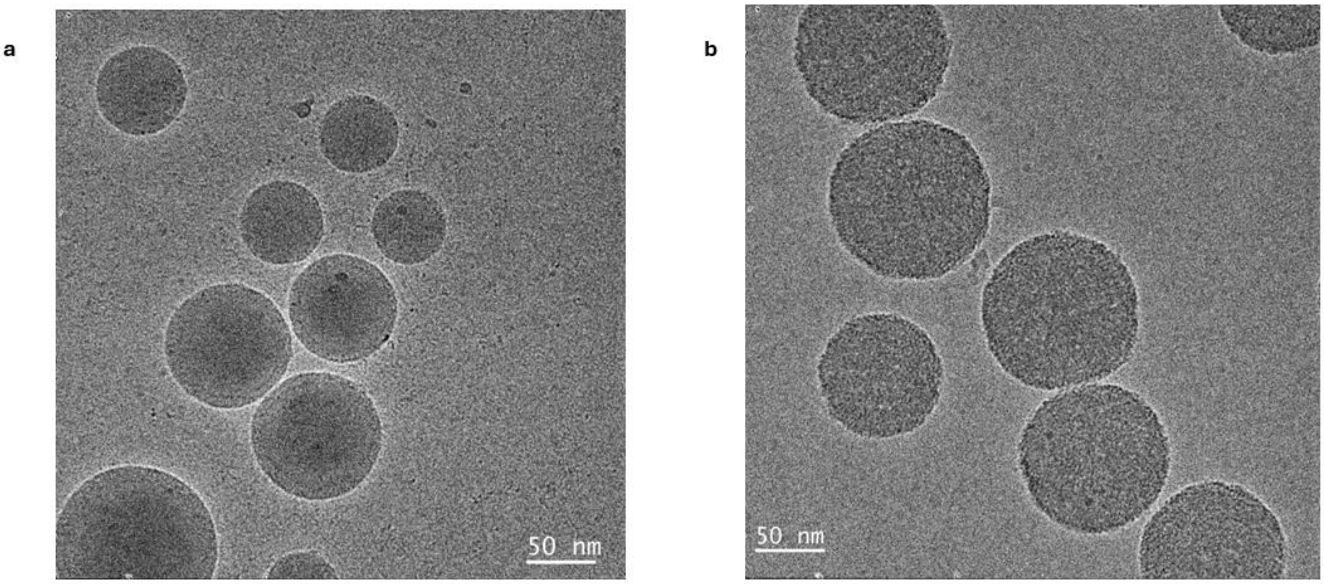


**Fig. S2. Representative Cryo-TEM images of PLGA (a) and i-Bind (b) NPs.**

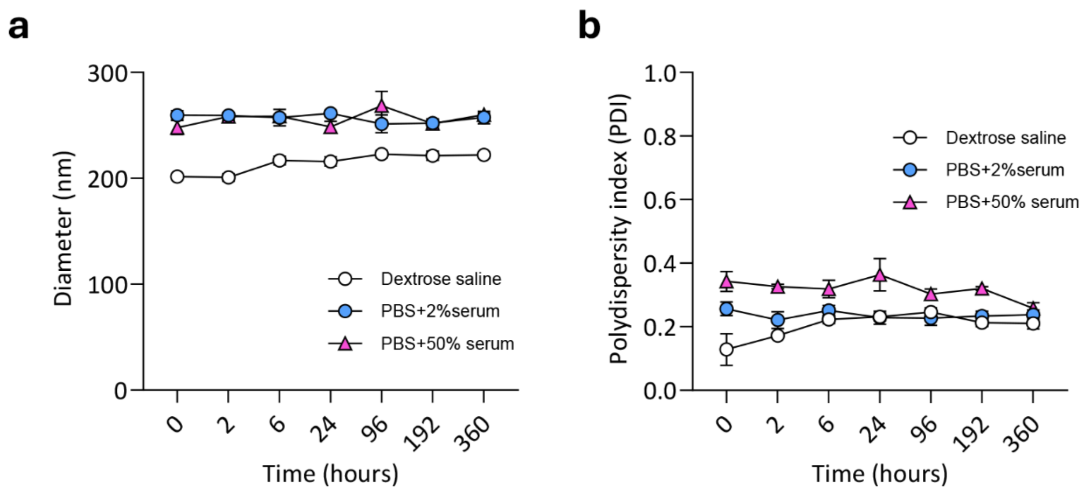


**Fig. S3. Colloidal stability of i-Bind NPs over 15 days. a**, Change of NP size. **b,** Change of NP polydispersity index. Data are presented as mean ± SEM.

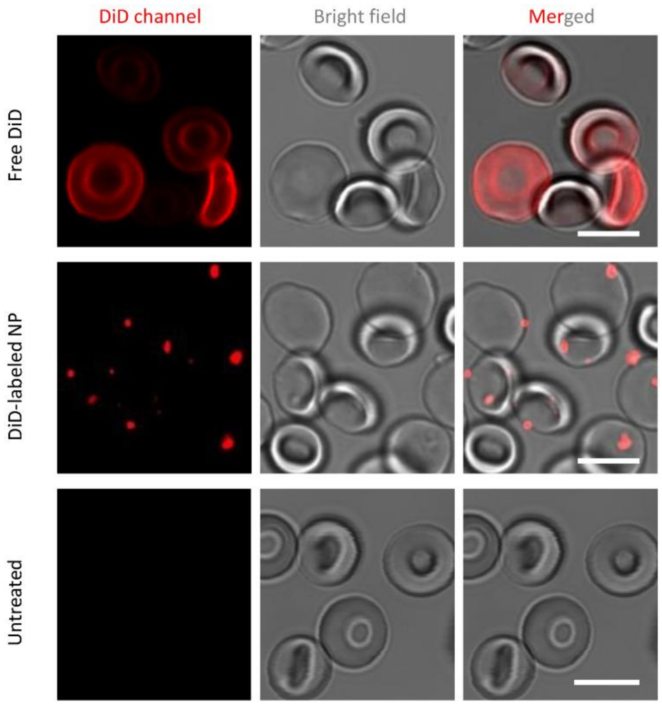


**Fig. S4. Evaluation of DiD dye transfer from DiD-labeled NPs to red blood cell membranes.** Free DiD uniformly stains RBC membranes and did not show similar particulate patterns as seen in DiD-labeled NP treated cells. Scale bars: 5μm.

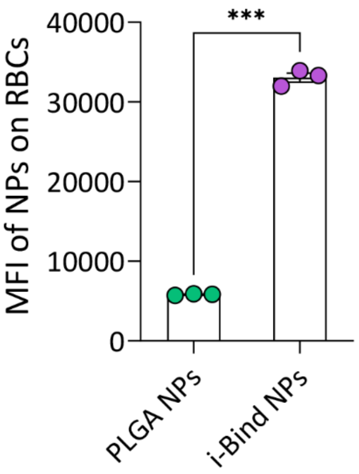


**Fig. S5. Number of NPs (indicated by MFI) bound to RBCs after free mixing in pure RBC samples (40% hematocrit).** Data are presented as mean ± SEM. Statistical analysis was conducted by two-tailed student's t test: *** $p < 0.001$.

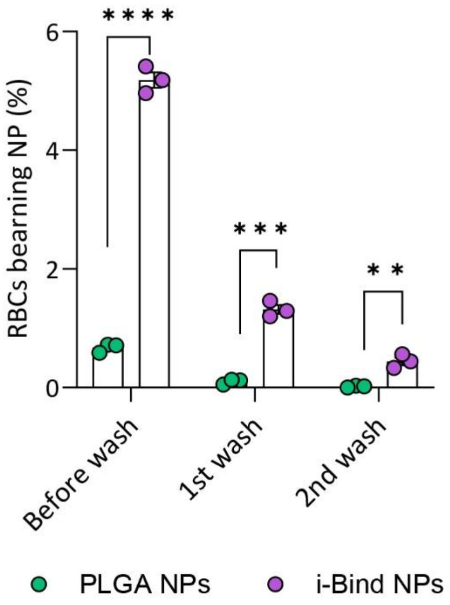


**Fig. S6. Percentage of RBCs carrying NPs when incubating NPs with mouse whole blood.** Data are presented as mean ± SEM. Statistical analysis was conducted by two-tailed student's t test: ** $p < 0.01$, *** $p < 0.001$, **** $p < 0.0001$.

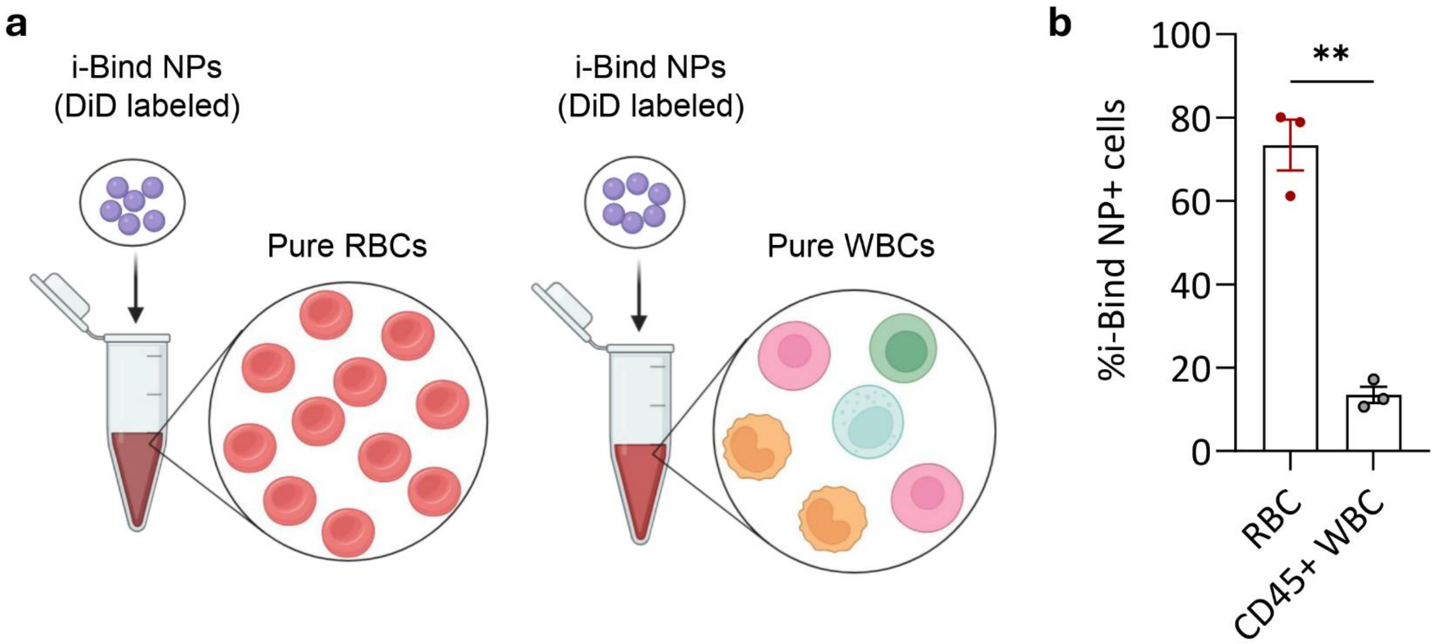


**Fig. S7. i-Bind NP hitchhiking using the same number of WBCs versus RBCs. a,** Schematic showing the experimental design to measure binding of DiD-labeled i-Bind NPs to RBCs or WBCs in pure RBC or WBC samples at same NP and cell concentration. Cells were incubated with NPs for 3 minutes. Created in BioRender. Zhao, Z. (2025) https://BioRender.com/q9rc2mv; https://BioRender.com/76otyy0. **b,** Percentage of NP-positive cells, RBC and WBC treated separately. Data are presented as mean ± SEM. Statistical analysis in (**b**) was conducted by two-tailed student's t test: ** $p < 0.01$.

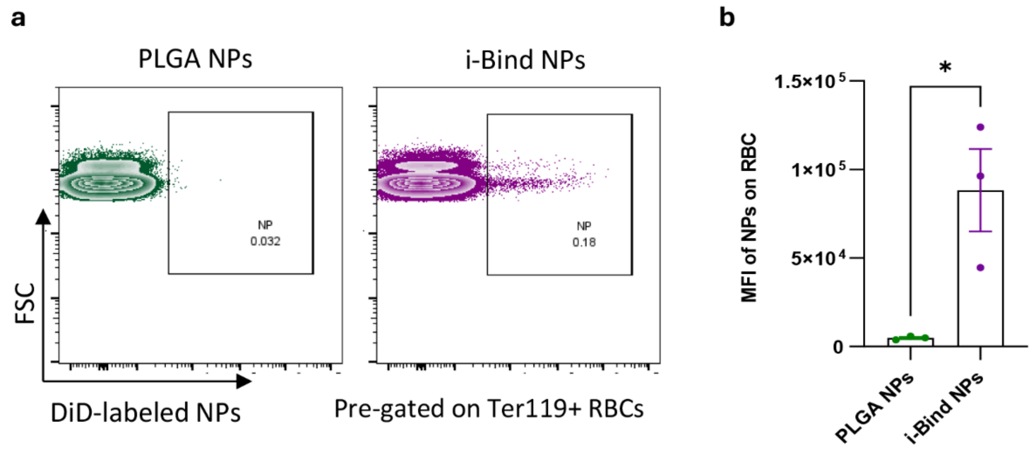


**Fig S8. *In vivo* NP association with RBCs 2 minutes after intravenous injection in healthy mice. a,** Representative flow plots showing NP binding to RBCs. **b,** Quantification of the number of NPs (indicated by MFI) bound to RBCs. Data in (**b**) are presented as mean ± SEM. Significantly different (two-tailed student's t test): * $p < 0.05$.

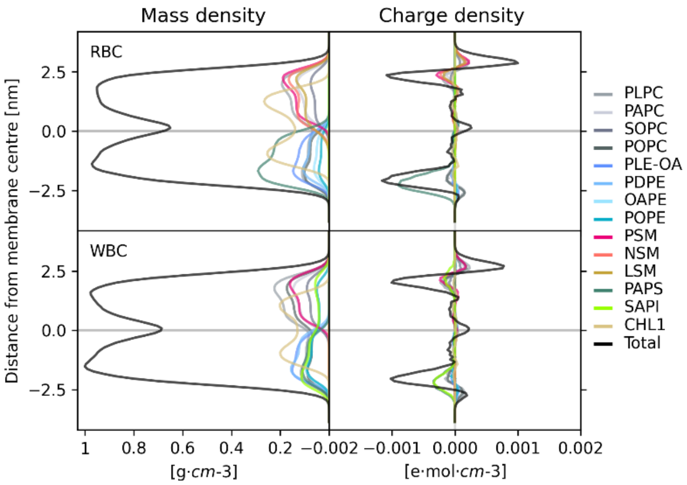


**Fig S9. The mass density and charge density profiles of the asymmetric membrane models as a function of the distance from the membrane center along the membrane normal.** 1-palmitoyl-2-linoleoyl-sn-glycero-3-phosphocholine (PLPC); 1-palmitoyl-2-arachidonoyl-sn-glycero-3-phosphocholine (PAPC); 1-stearoyl-2-oleoyl-sn-glycero-3-phosphocholine (SOPC); 1-palmitoyl-2-oleoyl-sn-glycero-3-phosphocholine (POPC); plasmalogen phosphatidylethanolamine (PLE-OA); 1-palmitoyl-2-docosahexaenoyl-sn-glycero-3-phosphoethanolamine (PDPE); 1-oleoyl-2-arachidonoyl-sn-glycero-3-phosphoethanolamine (OAPE); 1-palmitoyl-2-oleoyl-sn-glycero-3-phosphoethanolamine (POPE); 1-palmitoyl sphingomyelin (PSM); N-nervonoyl sphingomyelin (24:1 sphingomyelin) (NSM); lignoceroyl sphingomyelin (24:0 sphingomyelin) (LSM); 1-palmitoyl-2-arachidonoyl-sn-glycero-3-phospho-L-serine (PAPS); 1-stearoyl-2-arachidonoyl-sn-glycero-3-phospho-(1′-myo-inositol) (SAPI); cholesterol (CHL).

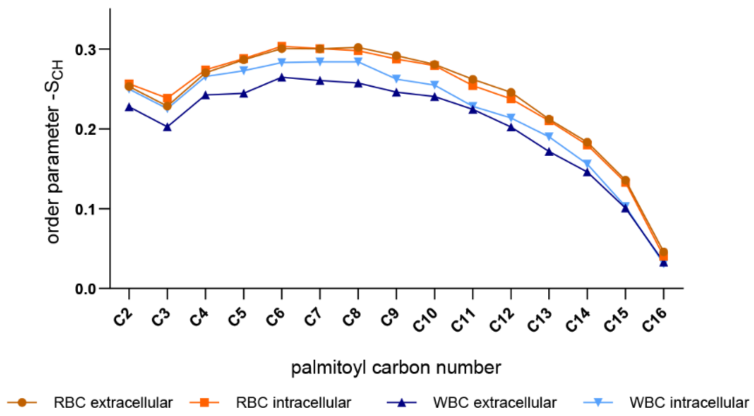


**Fig. S10. Order parameters for the two leaflets of asymmetric membranes, shown for the palmitoyl chains of PAPC (extracellular) or PAPS (intracellular) leaflets.** 1-palmitoyl-2-arachidonoyl-sn-glycero-3-phosphocholine (PAPC); 1-palmitoyl-2-arachidonoyl-sn-glycero-3-phospho-L-serine (PAPS).

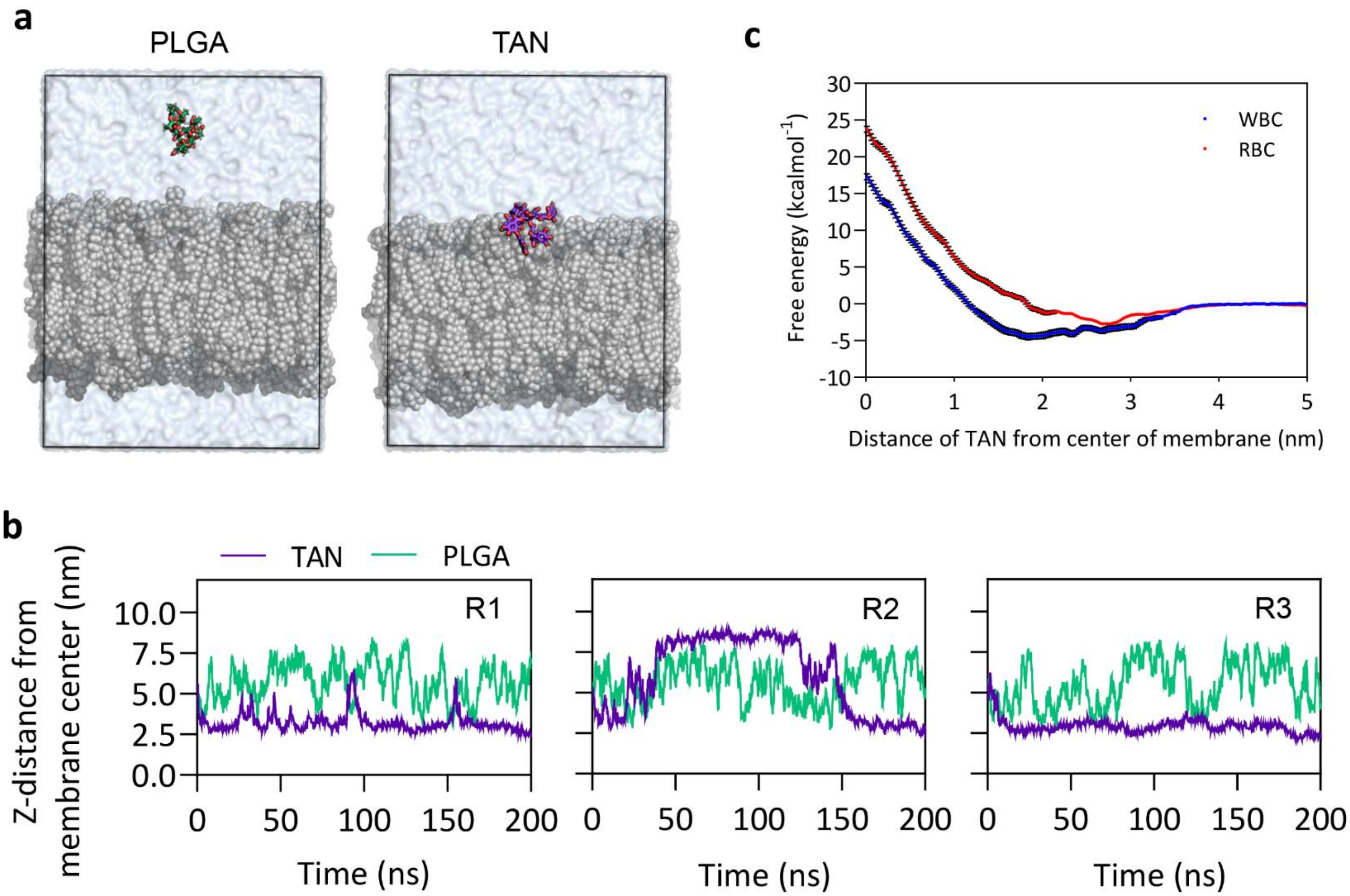


**Fig. S11. Simulation of PLGA or tannic acid (TAN) binding to WBC membrane. a,** Representative snapshots of the preferred position of PLGA and TAN with a WBC lipid membrane. TAN is depicted in purple sticks, PLGA in green, lipids in grey and water as blue surface. **b,** Time resolved distance evolution between the TAN and PLGA center of mass and the WBC membrane center of mass, projected to the z-axis. R1, R2, and R3 represent three replicates of simulations. **c,** Free energy profile of TAN with RBC and WBC membranes as a function of the TAN center of mass from the membrane center of mass projected on the membrane normal.

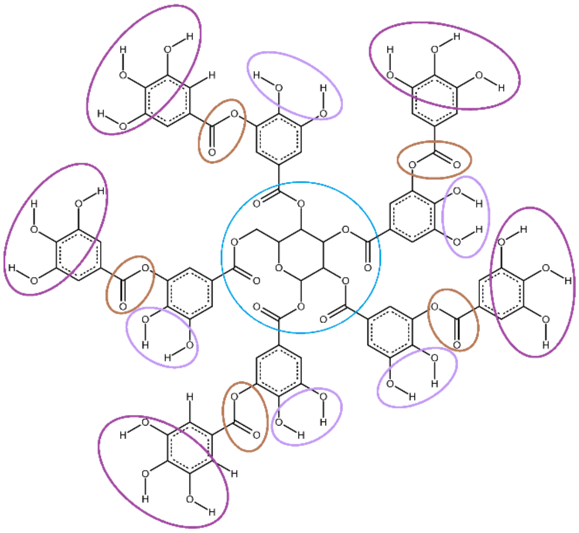

**Fig. S12. The structure of the tannic acid molecule with the defined layers of hydrogen bond-forming atoms.** Inner glucose layer in blue, middle gallic acid hydroxyls in light purple, ester layer in brown, outer gallic acid hydroxyl layer in dark purple.

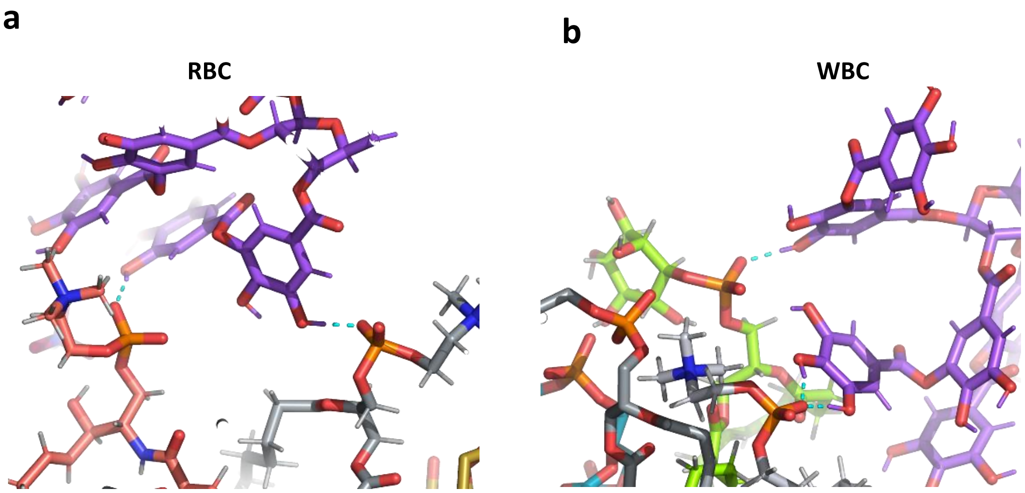


**Fig. S13. A detailed view of TAN (purple) forming hydrogen bonds (shown in cyan) with PLPC (light gray), NSM (pink), PAPC (dark gray) and SAPI (green). a,** RBC snapshot showing the middle and outer layers of the same TAN branch are involved in two hydrogen bonds with lipid molecules. **b,** WBC snapshot showing the middle and outer layers of gallic acid hydroxyls of different TAN branches are involved in three hydrogen bonds. 1-palmitoyl-2-linoleoyl-sn-glycero-3-phosphocholine (PLPC); N-nervonoyl sphingomyelin (24:1 sphingomyelin) (NSM); 1-palmitoyl-2-arachidonoyl-sn-glycero-3-phosphocholine (PAPC); 1-stearoyl-2-arachidonoyl-sn-glycero-3-phospho-(1′-myo-inositol) (SAPI).

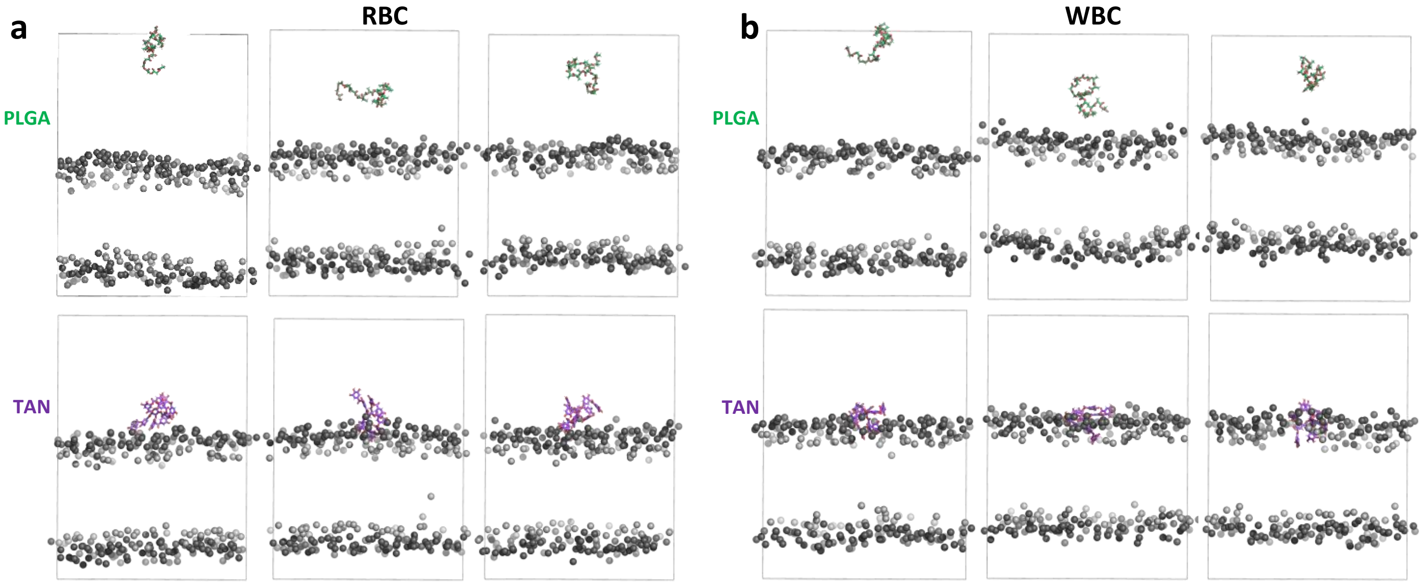


**Fig. S14. Final snapshots of the studied systems.** PLGA is depicted in green, and TAN in purple sticks; cholesterol oxygen atoms are light grey balls (the inner part of the membrane); headgroup nitrogens are dark green balls (the outer part of the membrane). The other atoms of the membrane, water, and ions are hidden for clarity. The simulation box is shown in lines.

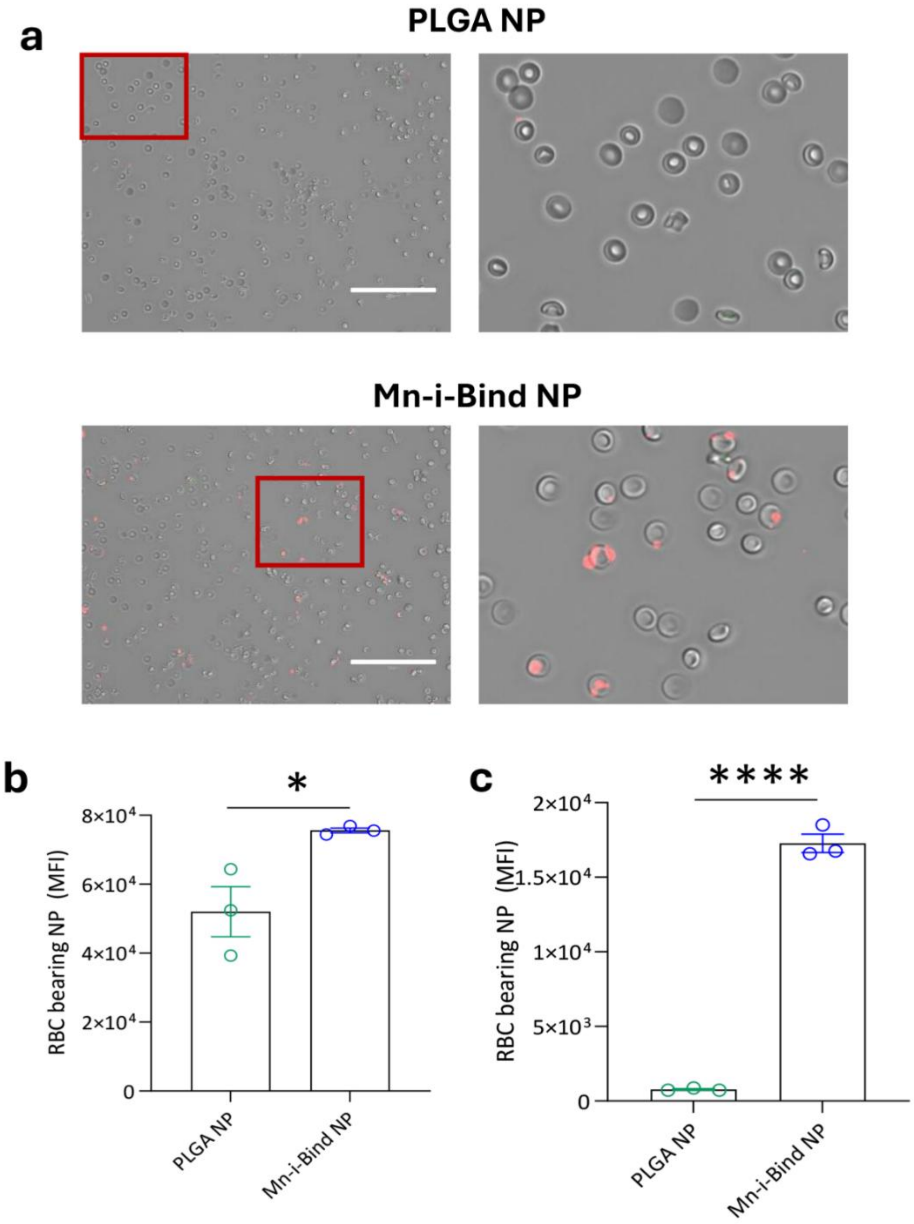


**Fig. S15. Mn-i-Bind NP hitchhiking to RBCs. a,** Fluorescent microscopic images overlayed on brightfield images, showing binding of DiD-labeled PLGA or Mn-i-Bind NPs to RBCs in pure RBC samples. Scale Bar: 75 μm. **b,** Number of NPs (indicated by MFI) bound to RBCs after free mixing in pure RBC samples. **c,** Number of NPs (indicated by MFI) bound to RBCs after free mixing in whole blood samples. Data in (**b-c**) are presented as mean ± SEM. Statistical analysis in (**b, c**) was conducted by two-tailed student's t test: * $p < 0.05$, **** $p < 0.0001$.

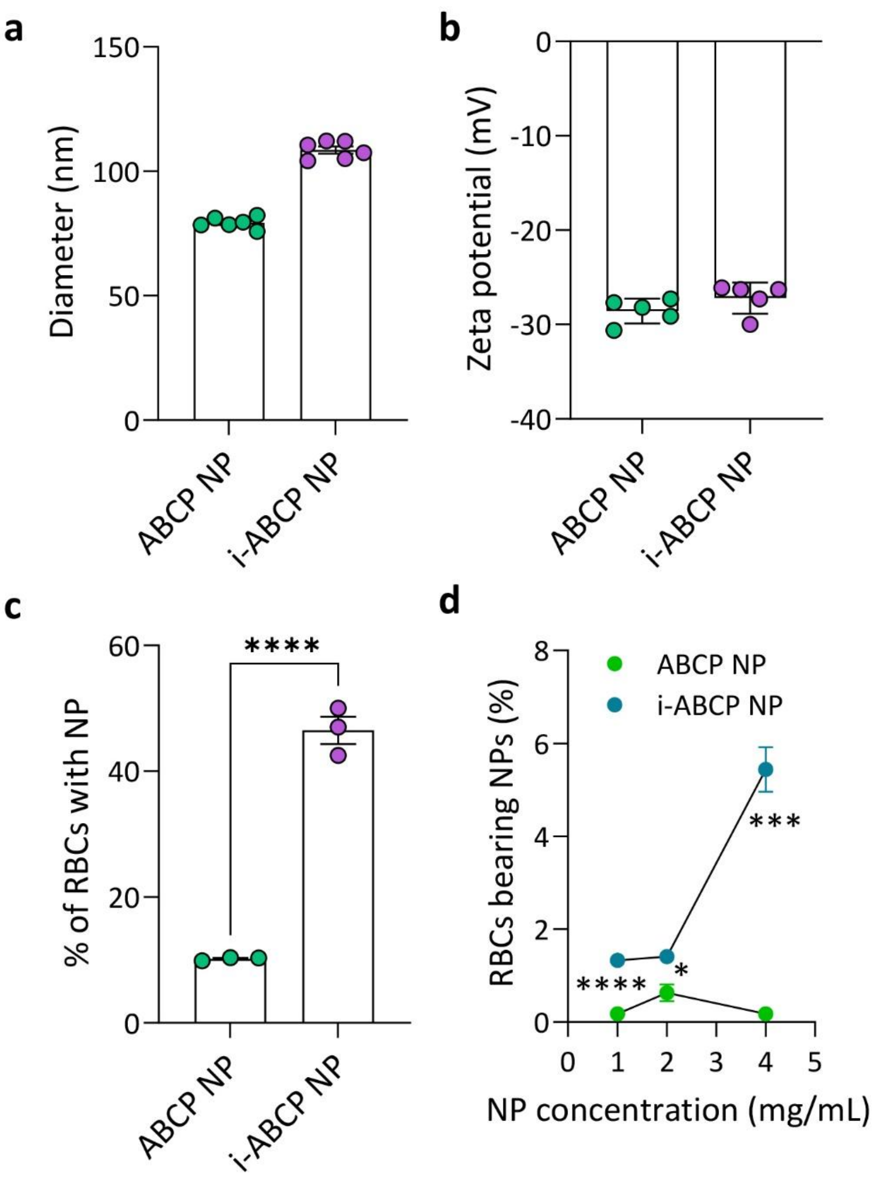


**Fig. S16. Characterization of the physicochemical properties of ABCP polymeric NPs and their binding to RBCs. a,** Diameter of NPs. **b,** Surface charge of NPs. **c,** Percentage of RBCs carrying NPs when incubating NPs with 10% purified RBCs. **d,** Percentage of RBCs carrying NPs when mixing NPs with mouse whole blood in the microfluid flow condition. Data in (**a-d**) are presented as mean ± SEM. Statistical analysis in (**c, d**) was conducted by two-tailed student's t test: * $p < 0.05$, *** $p < 0.001$, **** $p < 0.0001$.

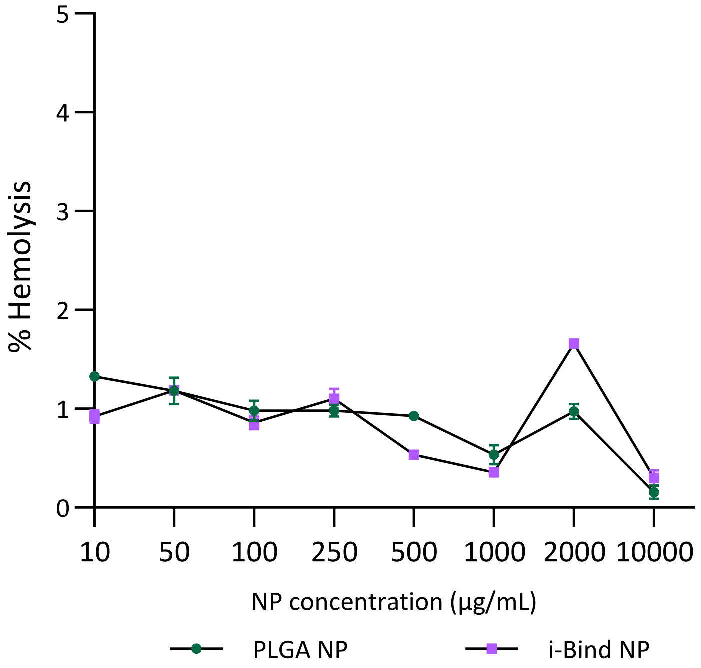


**Fig. S17. RBC hemolysis level induced by i-Bind NPs versus PLGA NPs at different NP concentrations.**

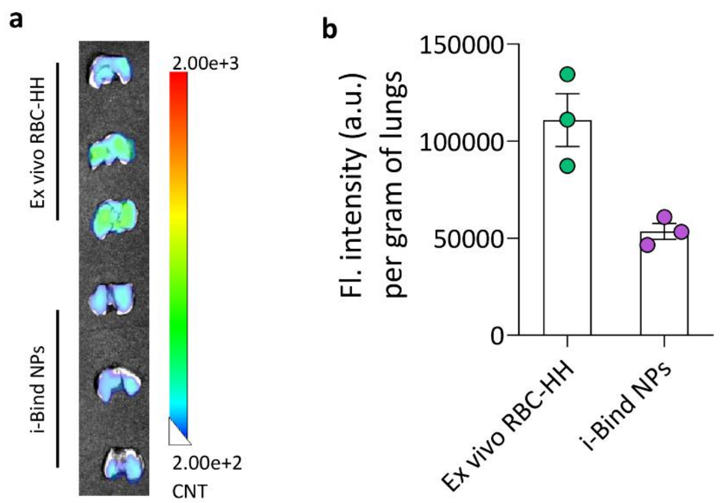


**Fig. S18. Biodistribution of i-Bind NPs versus *ex vivo* RBC-hitchhiked PLGA NPs 5 hours after intravenous administration in healthy mice. a,** Lago X fluorescence imaging data showing the relative accumulation of NPs in lungs. **b,** Quantification of the relative accumulation of NPs in lungs. Data in (**b**) are presented as mean ± SEM.

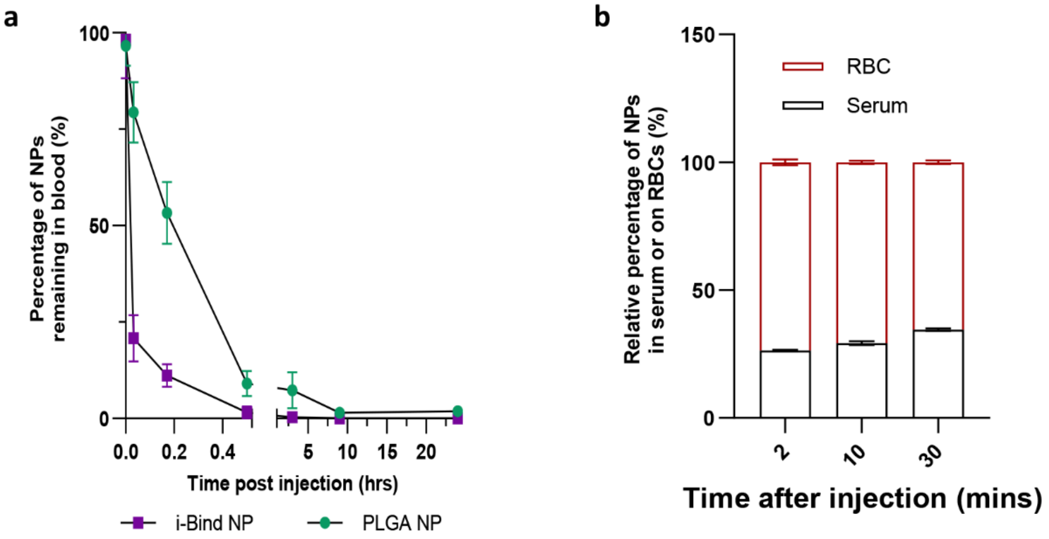


**Fig. S19. Pharmacokinetic study showing biodistribution of NPs in blood over 24 hours after intravenous injection in healthy mice. a,** Mouse 24 hours *in vivo* bloodstream circulation profile. Reading normalized to blood at 0 hour mixed with 100% dose of NPs (dilution was made based on the assumption that a mouse contains 80 mL/kg of blood). n=3 independent animals. **b,** Relative distribution of NPs remaining in circulation in serum and on RBCs at 2, 10, 30 minutes after administration. At each time point, the total NPs in serum and RBC fractions were counted as 100%. n=3 independent animals. Data in (**a-b**) are presented as mean ± SEM.

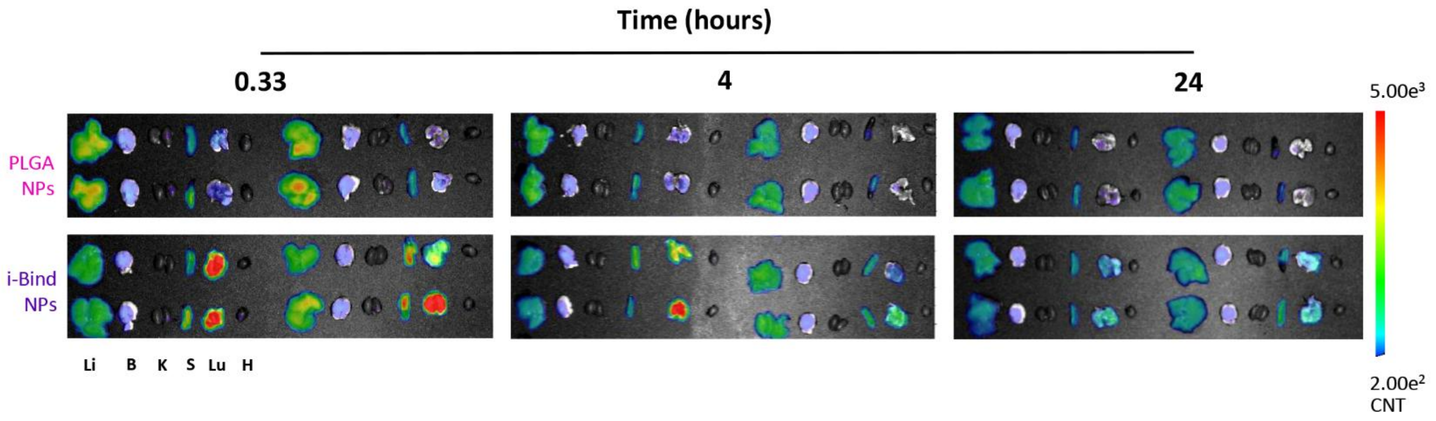


**Fig. S20. Lago X fluorescence images showing the biodistribution of NPs over 24 hours after intravenous injection in healthy mice.** Li, liver; B, brain; K, kidney; S, spleen; Lu, lung; H, heart.

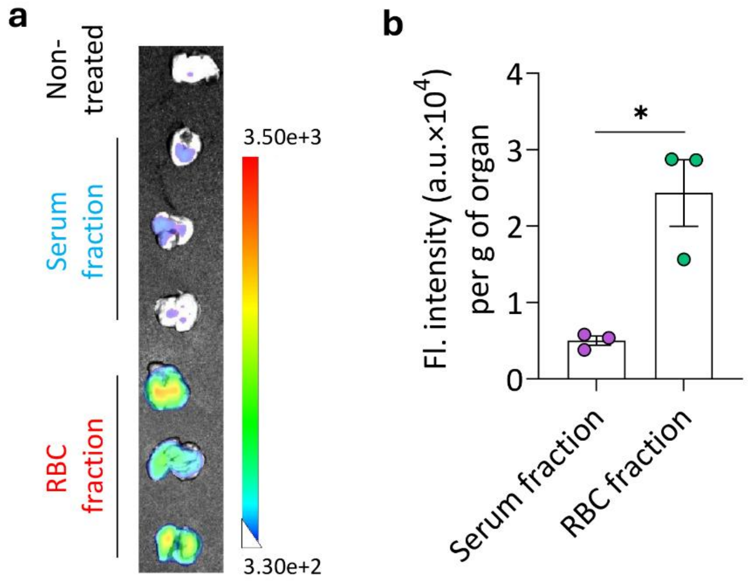


**Fig. S21. Evaluation of the relative contribution of serum and RBC fractions to i-Bind NP lung accumulation at equal NP doses.** The experimental design is similar to that shown in Fig. 3f, except that the serum and RBC fractions containing equal number of NPs were injected. **a,** Lago X image showing lung accumulation of i-Bind NPs 5 hours after intravenous injection of serum or RBC fractions containing same amount of NPs**. b,** Quantification of NP accumulation in the lungs. Data in (**b**) are presented as mean ± SEM. Significantly different (two-tailed student's t test): * $p < 0.05$.

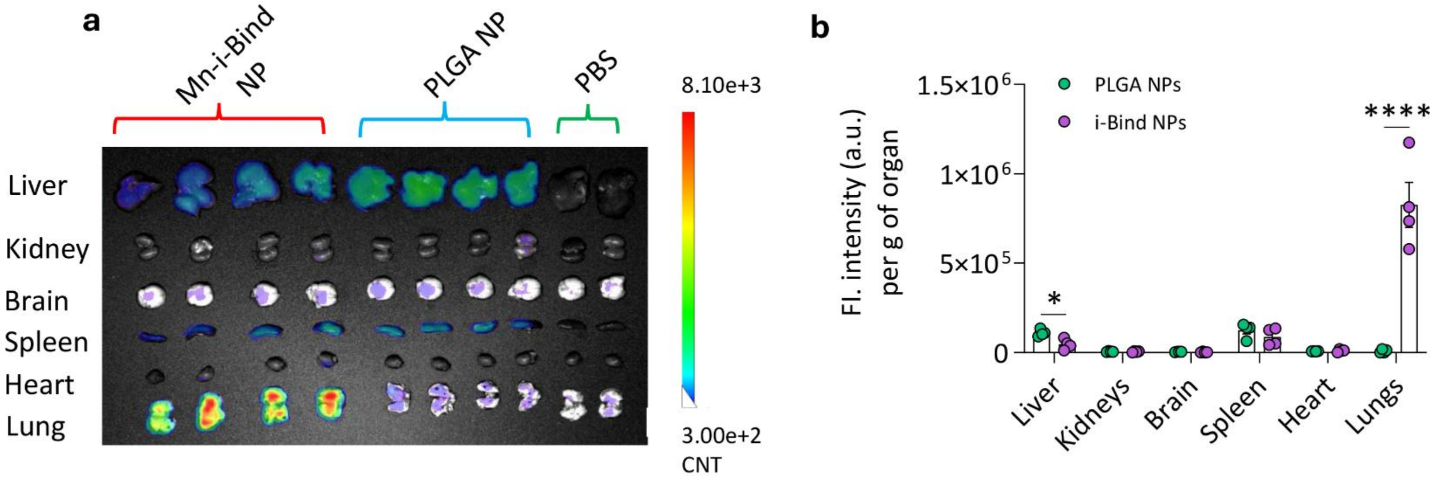


**Fig. S22. Biodistribution of i-Bind NPs prepared from Mn-TA complexation (Mn-i-Bind NPs) 5 hours after intravenous administration in healthy mice. a**, Lago X fluorescence imaging data showing the relative accumulation of NPs in major organs. **b,** Quantification of the relative accumulation of NPs in different organs. Data in (**b**) are presented as mean ± SEM. Significantly different (two-tailed student's t test): * $p < 0.05$, **** $p < 0.0001$.

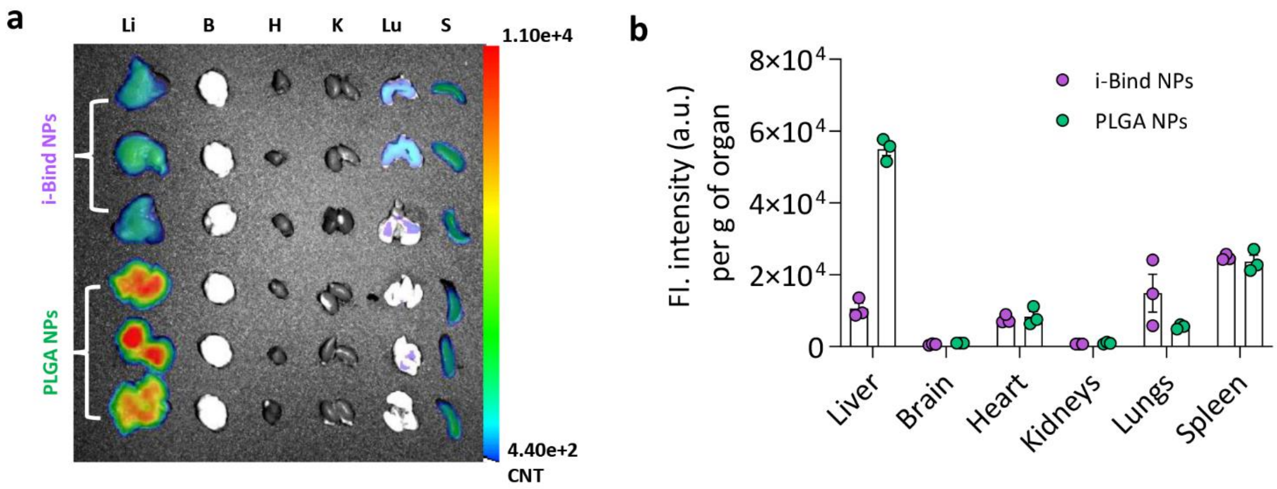


**Fig. S23. Biodistribution of NPs 5 hours after intravenous administration in the LPS inhalation-induced ALI model. a**, Lago X fluorescence imaging data showing the relative accumulation of NPs in major organs. Li, liver; B, brain; K, kidney; S, spleen; Lu, lung; H, heart. **b,** Quantification of the relative accumulation of NPs in different organs. Data in (**b**) are presented as mean ± SEM.

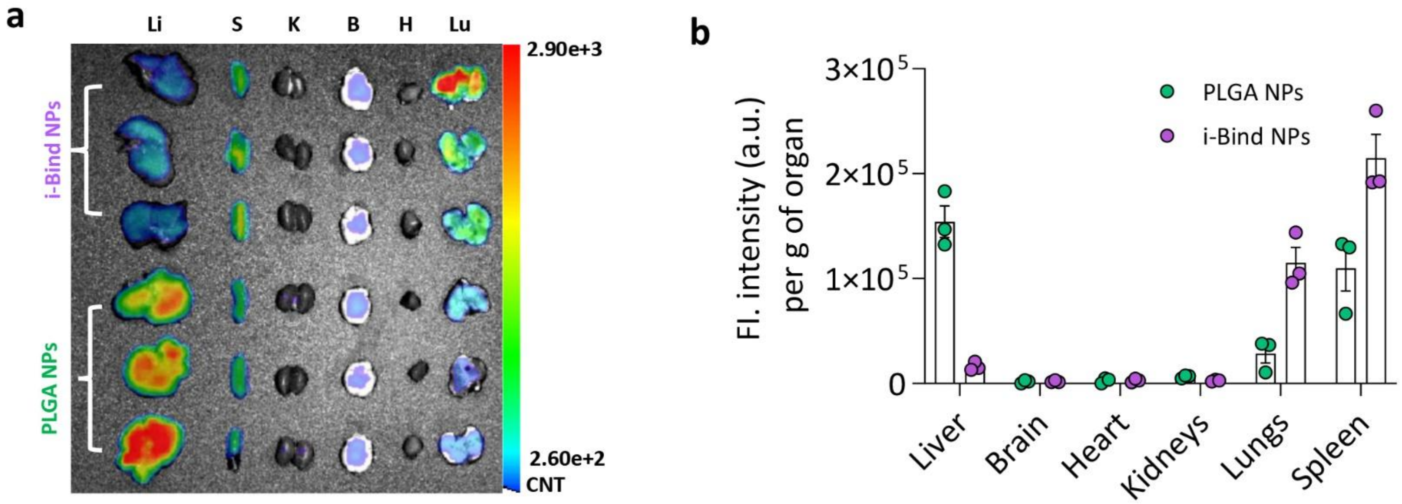


**Fig. S24. Biodistribution of NPs 5 hours after intravenous administration in the B16F10-Luc melanoma lung metastasis model. a,** Lago X fluorescence imaging data showing the relative accumulation of NPs in major organs. Li, liver; B, brain; K, kidney; S, spleen; Lu, lung; H, heart. **b,** Quantification of the relative accumulation of NPs in different organs. Data in (**b**) are presented as mean ± SEM.

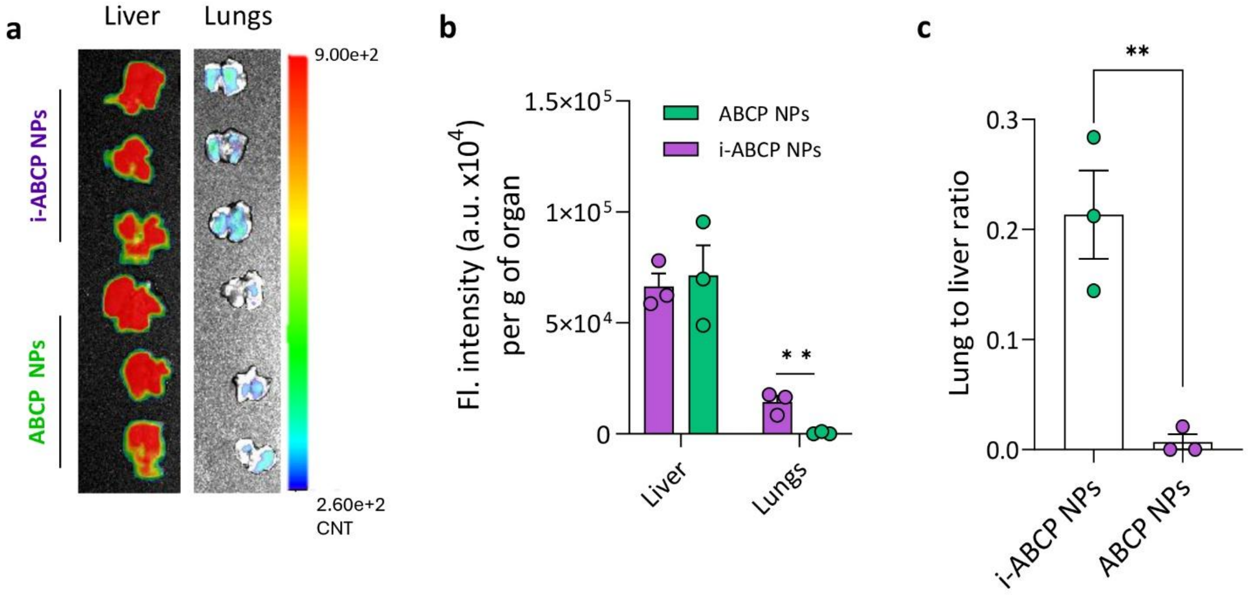


**Fig. S25. Biodistribution of non-coated or coated ABCP NPs 5 hours after intravenous administration in healthy mice. a,** Lago X fluorescence imaging data showing the relative accumulation of NPs in liver and lungs. **b,** Quantification of the relative accumulation of NPs in liver and lungs. **c,** Lung to liver ratio of NP accumulation. Data in **(b, c)** are presented as mean ± SEM. Statistical analysis in **(b, c)** was conducted by two-tailed student's t test: ** $p < 0.01$.

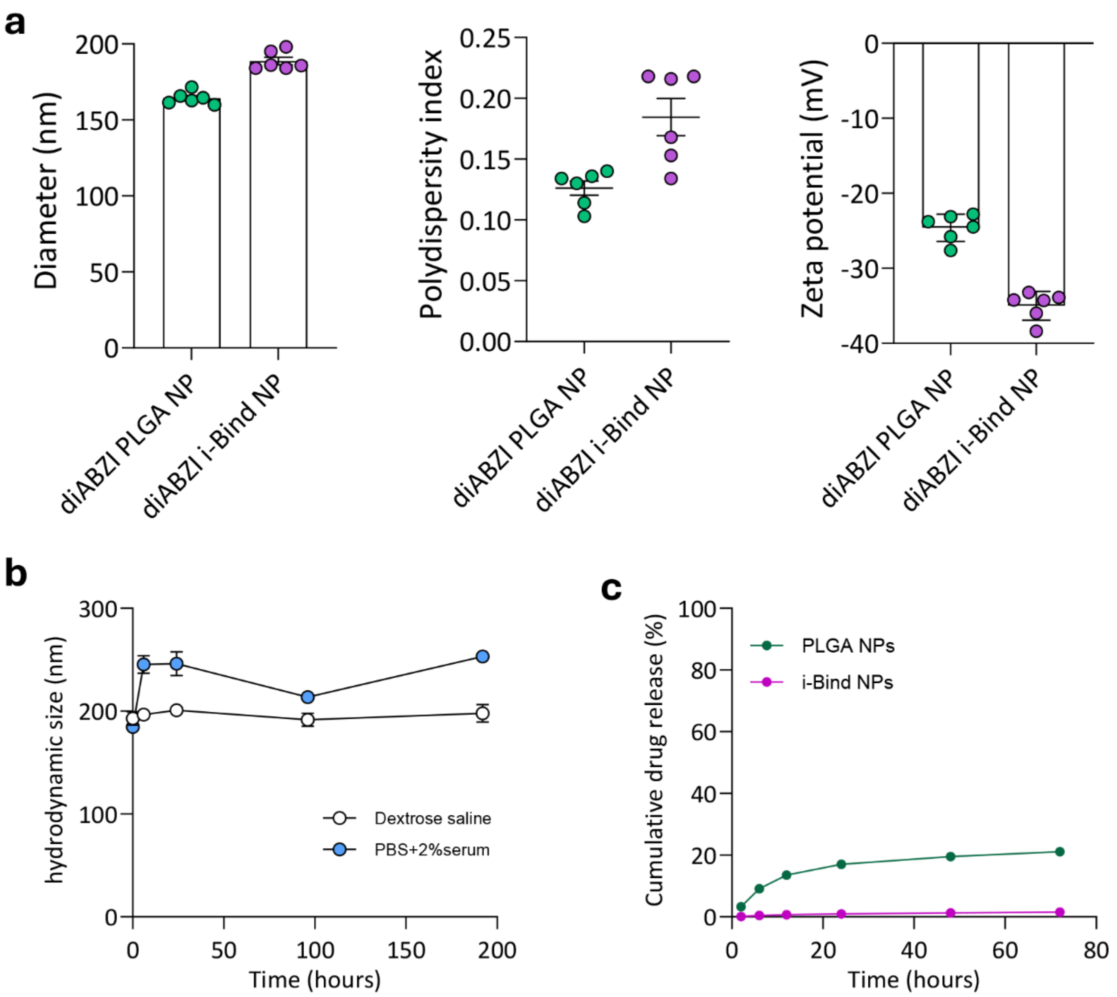


**Fig. S26. Characterization of diABZI-loaded i-Bind NPs. a,** Physicochemical characterization showing hydrodynamic size, polydispersity index, and zeta potential of diABZI-loaded i-Bind NPs. **b,** Colloidal stability of diABZI i-Bind NPs in different buffers. **c,** Drug release profile of diABZI from NPs. Release study was conducted in PBS (10 mM, pH 7.4), substituted with 1% FBS at 37 °C. n=3 independent samples for each NP type. Data in **(a-c)** are presented as mean ± SEM.

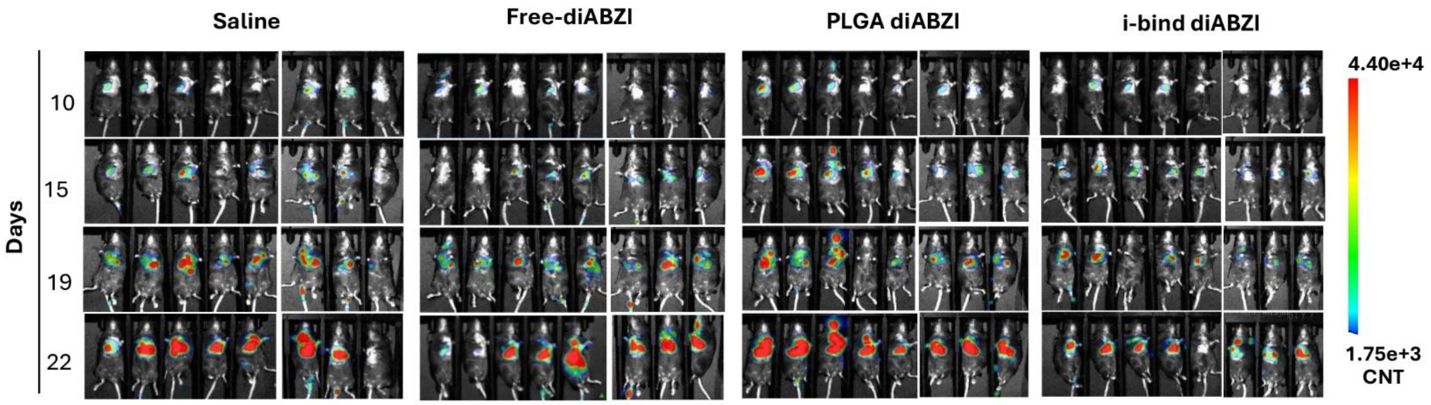


**Fig. S27. Lago X bioluminescence imaging data showing the progression of lung metastases in mice treated with different diABZI formulations.** Note: Representative mouse images were shown in Fig. 5b with a different scale range.

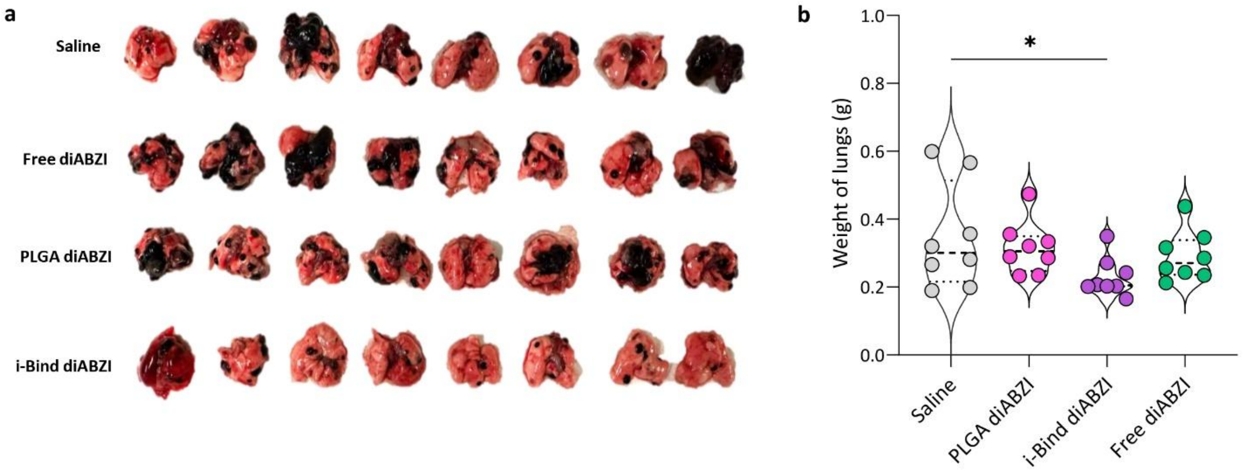


**Fig. S28. Analysis of the lungs of mice following different treatments on day 23 in the B16F10-Luc lung metastasis model. a,** Images of lungs. **b,** Weight of lungs. Data in (**b**) are presented as mean ± SEM. Statistical analysis in (**b**) was conducted by one-way ANOVA: * $p < 0.05$.

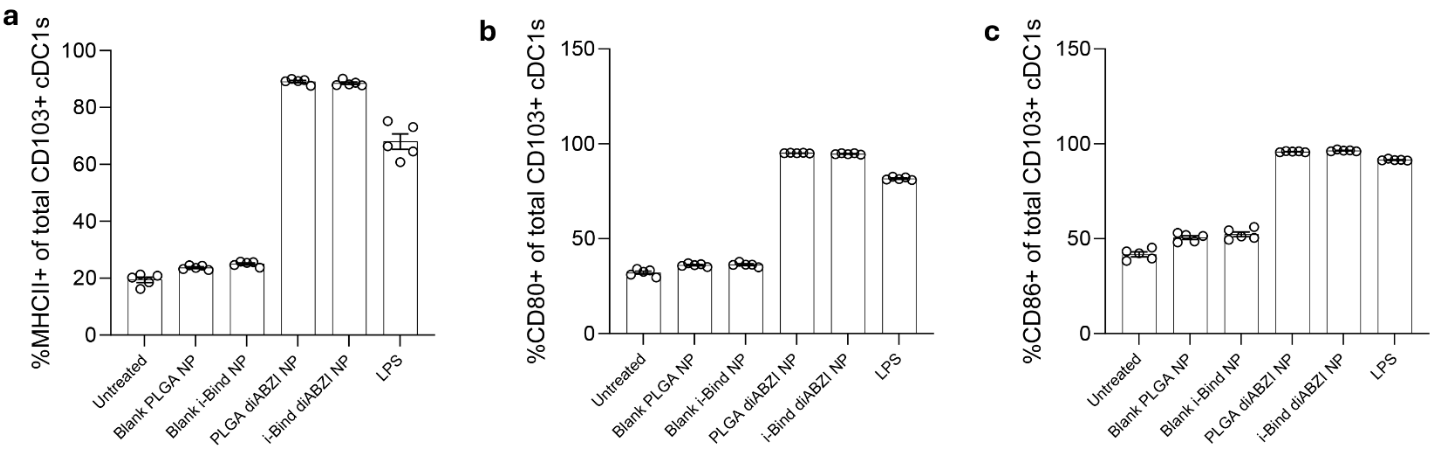


**Fig. S29. Activation of DCs (CD103$^+$ cDC1s) by blank or diABZI-loaded i-Bind NPs after 12-hour incubation. a-c,** Percentage of DCs positive for activation markers MHCII (**a**), CD80 (**b**), and CD86 (**c**). Data in (**a-c**) are presented as mean ± SEM.

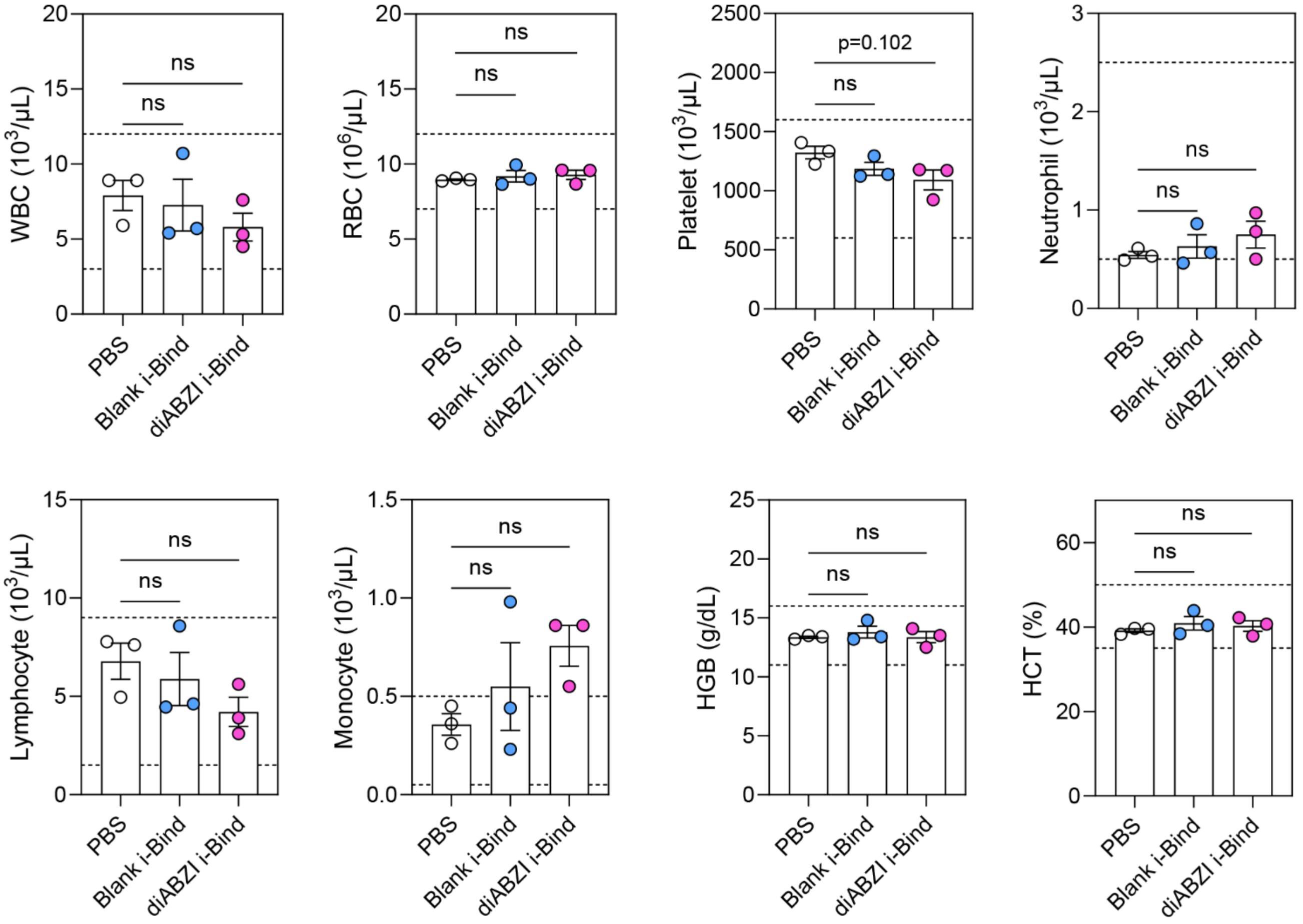


**Fig. S30. Hematological analysis of mice after intravenous administration of blank or diABZI-loaded i-Bind NPs.** Hematological parameters were measured on day 7 post-administration of NPs. (n=3 biological independent mice). Data presented as mean ± SEM. ns: no statistically significant difference (one-way ANOVA followed by Dunnett's test). Dashed lines indicate normal range. WBC: white blood cell; RBC: red blood cell; HGB: hemoglobin; HCT: hematocrit.

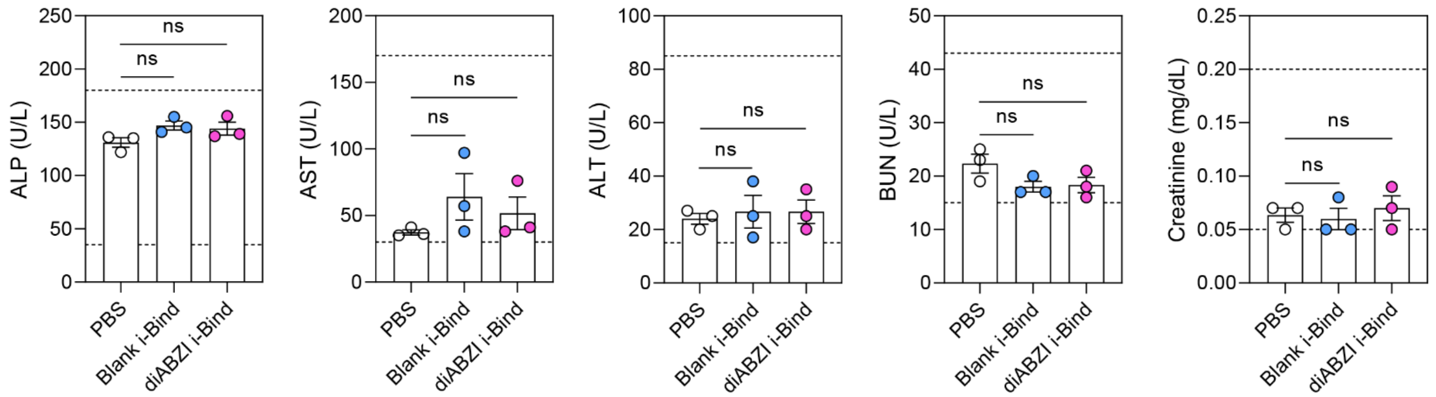


**Fig. S31. Serum chemistry of mice after intravenous administration of i-Bind NPs.** Serum chemistry parameters were measured 7 days post-intravenous administration of NPs (n=3 biological independent mice). Data presented as mean ± SEM. ns: no statistically significant difference (one-way ANOVA followed by Dunnett's test). Dashed lines indicate normal range. ALP: alkaline phosphatase; AST: aspartate aminotransferase; ALT: alanine transaminase; BUN: blood urea nitrogen.

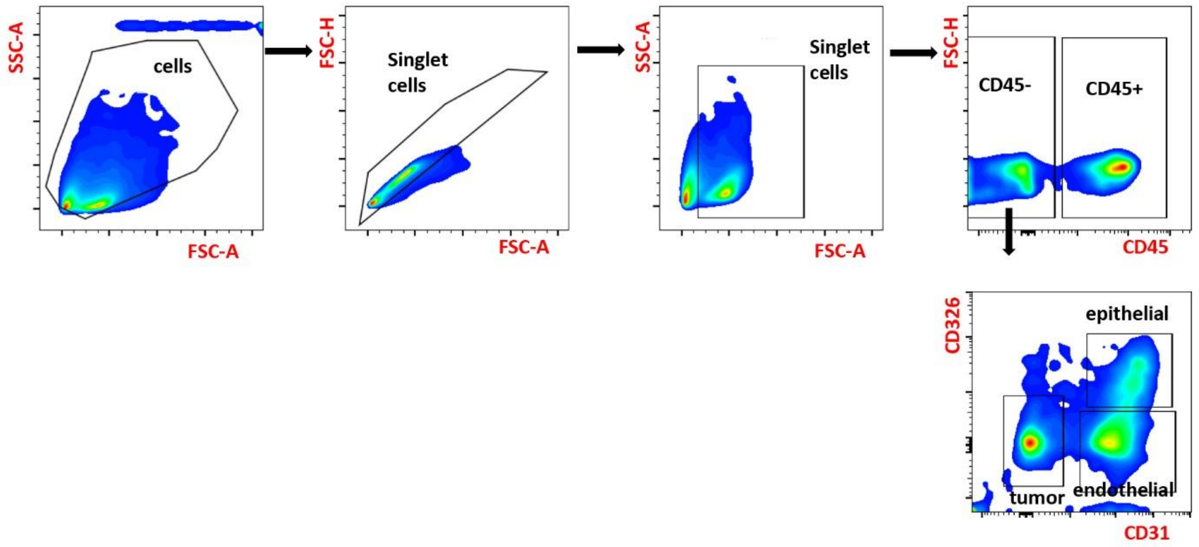


**Fig. S32. Representative gating strategy for identification of endothelial, epithelial, and/or tumor cells in the lung for the biodistribution study.**

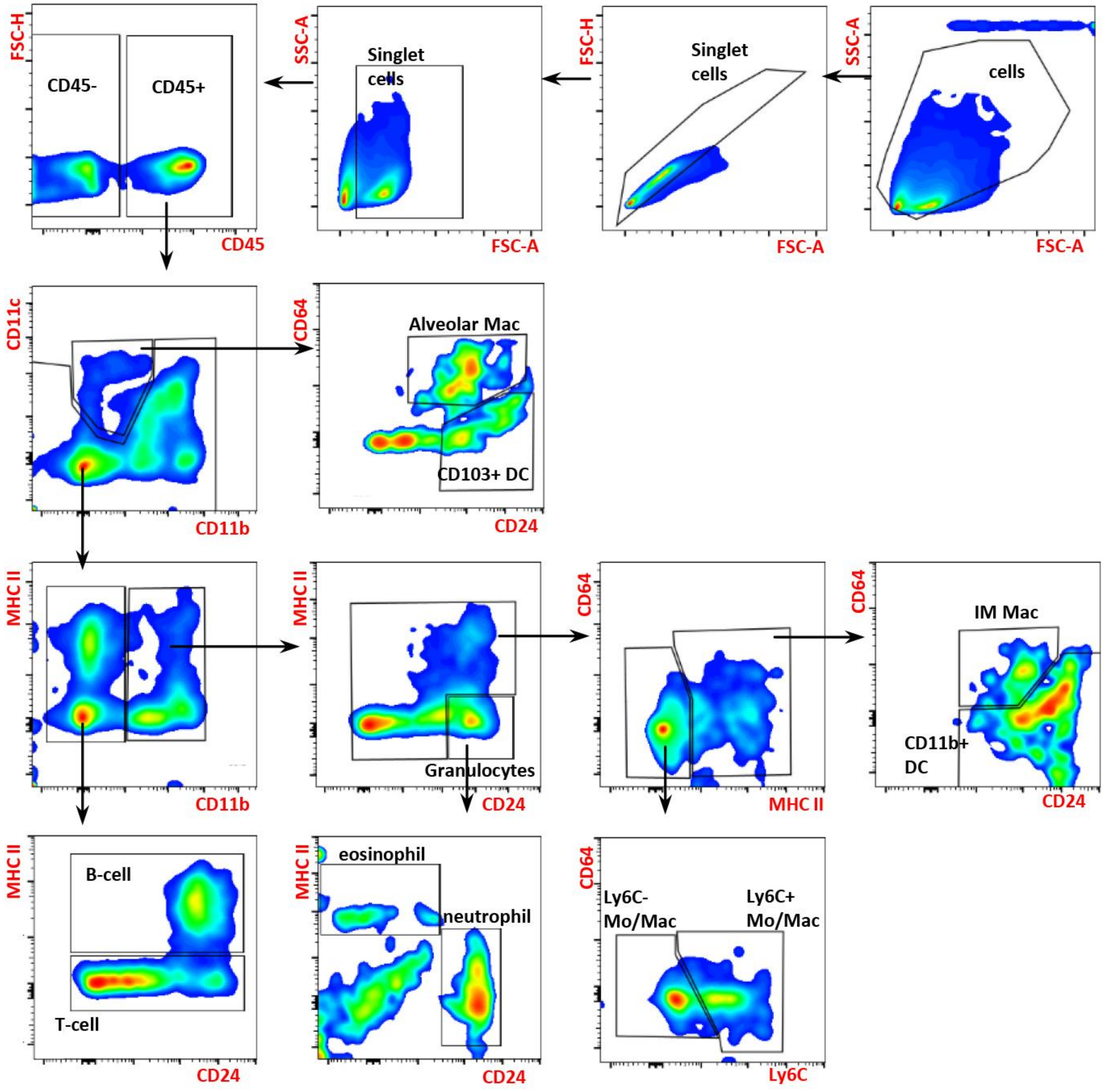


**Fig. S33. Representative gating strategy for the identification of pulmonary immune cell populations for the biodistribution study.**

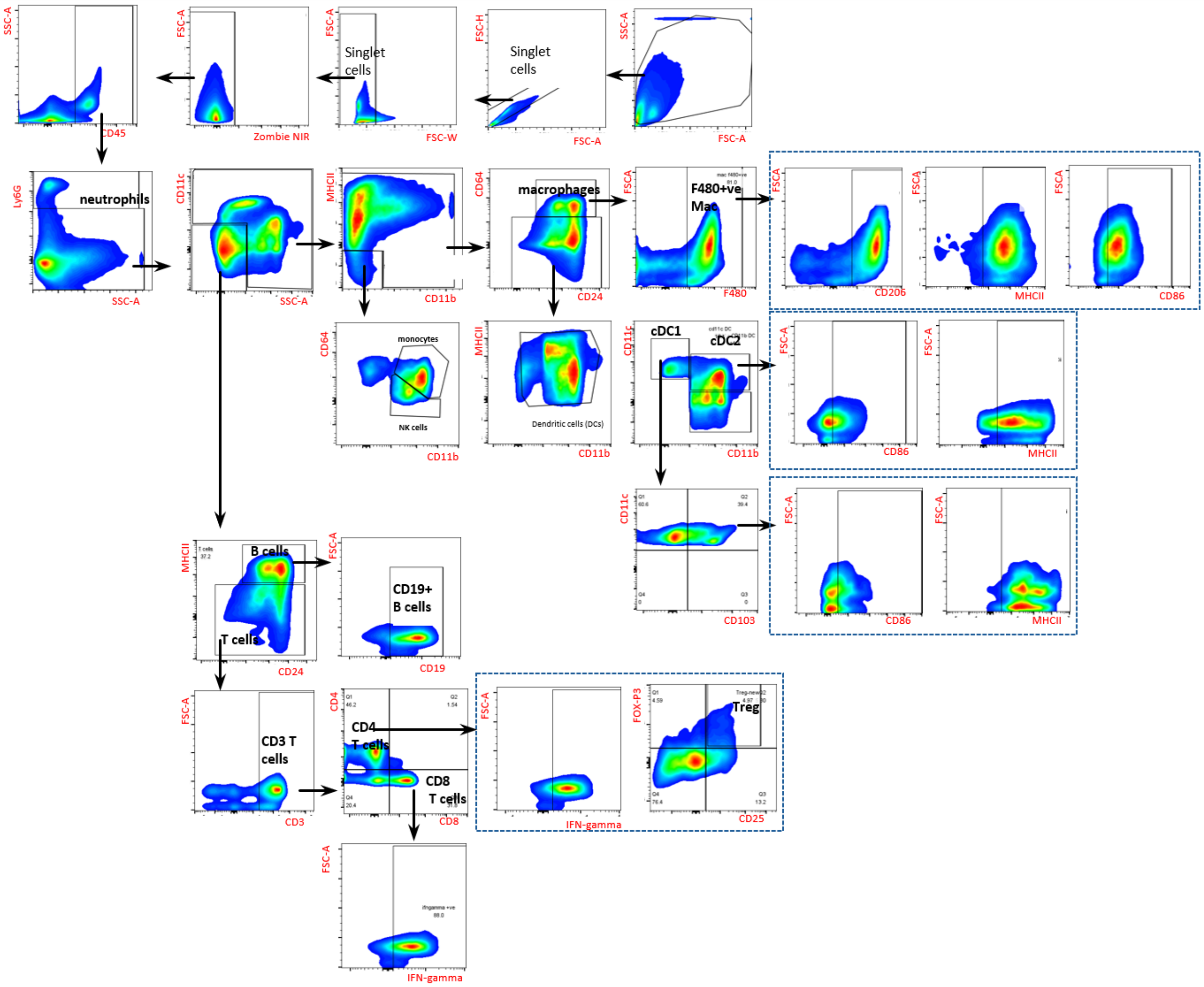


**Fig. S34. Representative gating strategy for the identification of pulmonary immune cell populations for the therapeutic efficacy study.**

**Table S1.**
**Physicochemical properties of polyphenol-coated nanoparticles.**

| | Average Size (nm) | Polydispersity index | Zeta Potential (mV) |
|---|---|---|---|
| PLGA NP (non-coated) | 168.7 ± 1.7 | 0.021 ± 0.013 | -18.54 ± 0.69 |
| Caffeic acid-coated NP (CA NP) | 172.3 ± 2.3 | 0.026 ± 0.014 | -18.90 ± 1.06 |
| Catechin-coated NP (CT NP) | 168.4 ± 1.0 | 0.006 ± 0.003 | -19.34 ± 1.17 |
| Resveratrol-coated NP (RV NP) | 169.5 ± 1.2 | 0.031 ± 0.023 | -10.80 ± 3.52 |
| Epigallocatechin-coated NP (ECG NP) | 156.8 ± 1.7 | 0.046 ± 0.038 | +0.41 ± 0.95 |
| Gallic acid-coated NP (GA NP) | 192.9 ± 5.8 | 0.046 ± 0.027 | -18.90 ± 1.36 |
| Tannic acid-coated NP (TA NP) | 198.2 ± 6.5 | 0.085 ± 0.051 | -34.06 ± 2.27 |

**Table S2.**
**The number of lipids in each leaflet of the studied systems. PE-O denotes plasmogen lipids.**

| Molecule | | | RBC (# of lipids) | | WBC (# of lipids) | |
|---|---|---|---|---|---|---|
| Residue | Tail | head | extracellular | intracellular | extracellular | intracellular |
| PLPC | 16:0–18:2 | PC | 23 | 13 | 27 | 14 |
| PAPC | 16:0–20:4 | PC | 15 | | 17 | |
| SOPC | 18:0–18:1 | PC | 9 | | 11 | |
| POPC | 16:0–18:1 | PC | | 12 | 21 | 11 |
| PLEOA | 18:1–20:4 | PE-O | | 17 | | 18 |
| PDoPE | 16:0–22:6 | PE | | 12 | | 18 |
| OAPE | 18:1–20:4 | PE | | 7 | | 11 |
| POPE | 16:0–18:1 | PE | | 5 | 6 | 11 |
| PSM | 18:1-16:0 | SM | 23 | | 20 | |
| NSM | 18:1-24:1 | SM | 20 | | | |
| LSM | 18:1-24:0 | SM | 16 | | | |
| PAPS | 16:0–20:4 | PS | | 34 | | 13 |
| SAPI | 18:0–20:4 | PI | | | 6 | 12 |
| CHL | - | - | 42 | 40 | 32 | 32 |

**Abbreviations:** 1-palmitoyl-2-linoleoyl-sn-glycero-3-phosphocholine (PLPC); 1-palmitoyl-2-arachidonoyl-sn-glycero-3-phosphocholine (PAPC); 1-stearoyl-2-oleoyl-sn-glycero-3-phosphocholine (SOPC); 1-palmitoyl-2-oleoyl-sn-glycero-3-phosphocholine (POPC); plasmalogen phosphatidylethanolamine (PLEOA);
1-palmitoyl-2-docosahexaenoyl-sn-glycero-3-phosphoethanolamine (PDoPE); 1-oleoyl-2-arachidonoyl-sn-glycero-3-phosphoethanolamine (OAPE); 1-palmitoyl-2-oleoyl-sn-glycero-3-phosphoethanolamine (POPE); 1-palmitoyl sphingomyelin (PSM); N-nervonoyl sphingomyelin (24:1 sphingomyelin) (NSM); lignoceroyl sphingomyelin (24:0 sphingomyelin) (LSM); 1-palmitoyl-2-arachidonoyl-sn-glycero-3-phospho-L-serine (PAPS); 1-stearoyl-2-arachidonoyl-sn-glycero-3-phospho-(1′-myo-inositol) (SAPI); cholesterol (CHL).

**Table S3.**
**Average number of hydrogen bonds per frame of PLGA/TAN with water or membrane, number of intramolecular hydrogen bonds and the average of a total number of hydrogen bonds formed by TAN or PLGA calculated over the last 50 ns of the three replicas.**

| | # of hydrogen bonds | | | |
|---|---|---|---|---|
| | RBC-PLGA | RBC-TAN | WBC-PLGA | WBC-TAN |
| Water | 18 ± 2.3 | 21.7 ± 1 | 17.9 ± 0.2 | 21.5 ± 2.3 |
| Membrane | 0 ± 1.4 | 2.8 ± 0.3 | 0 ± 0 | 4.1 ± 1.4 |
| Intramolecular | 0 ± 0 | 0.3 ± 0.1 | 0 ± 0 | 0.2 ± 0 |
| Total | 18.1 ± 1 | 24.7 ± 0.7 | 17.9 ± 0.2 | 25.8 ± 1 |

**Table S4.**
**Average number of hydrogen bonds per frame between atoms of the respective layer and water or membrane is shown.**

| | # of hydrogen bonds | | | |
|---|---|---|---|---|
| | RBC | WBC | RBC | WBC |
| | water | | membrane | |
| Inner | 0.7 ± 0.1 | 0.6 ± 0.1 | 0 ± 0 | 0 ± 0 |
| Middle | 7.7 ± 0.9 | 8.1 ± 1.1 | 1.2 ± 0.8 | 1.3 ± 1 |
| Ester | 3.5 ± 0.1 | 3.6 ± 0.1 | 0 ± 0 | 0 ± 0 |
| Outer | 9.8 ± 0.8 | 9.1 ± 1.2 | 1.6 ± 0.6 | 2.8 ± 0.4 |

**Table S5.**
**Number of hydrogen bonds between TAN and lipid atoms, by the atom type.**

| CHARMM36 atom type | Atom description | # of H-bonds | |
|---|---|---|---|
| | | RBC | WBC |
| O2L | Phospholipid phosphate =O | 2.4 ± 0.6 | 3.3 ± 0.7 |
| OBL | Phospholipid glycerol ester oxygen | 0 ± 0 | 0.4 ± 0.3 |
| O | Sphingomyelin sidechain ester | 0.1 ± 0.3 | 0.2 ± 0.3 |
| OHL | Sphingosine chain/cholesterol hydroxyl | 0.2 ± 0.3 | 0 ± 0 |
| OSLP | Phospholipid phosphate -O- | 0 ± 0 | 0.1 ± 0 |
| OC311 | Inositol ring hydroxyl | 0 ± 0 | 0.1 ± 0 |
| NHL | Sphingosine chain amine | 0 ± 0 | 0 ± 0.1 |

O2L: Phospholipid phosphate double-bond oxygen (P=O)
OBL: Phospholipid glycerol ester oxygen
O: Sphingomyelin ester/carbonyl oxygen
OHL: Sphingosine-chain or cholesterol hydroxyl oxygen
OSLP: Phospholipid phosphate single-bond oxygen
OC311: Inositol-ring hydroxyl oxygen
NHL: Sphingosine-chain amine nitrogen
